# Inference of Genomic Landscapes using Ordered Hidden Markov Models with Emission Densities (oHMMed)

**DOI:** 10.1101/2023.06.26.546495

**Authors:** Claus Vogl, Mariia Karapetiants, Burçin Yıldırım, Hrönn Kjartansdóttir, Carolin Kosiol, Juraj Bergman, Michal Majka, Lynette Caitlin Mikula

## Abstract

**Background:** Genomes are inherently inhomogeneous, with features such as base composition, recombination, gene density, and gene expression varying along chromosomes. Evolutionary, biological, and biomedical analyses aim to quantify this variation, account for it during inference procedures, and ultimately determine the causal processes behind it. Since sequential observations along chromosomes are not independent, it is unsurprising that autocorrelation patterns have been observed *e.g.,* in human base composition.

In this article, we develop a class of Hidden Markov Models (HMMs) called oHMMed (ordered HMM with emission densities, the corresponding R package of the same name is available on CRAN): They identify the number of comparably homogeneous regions within autocorrelated observed sequences. These are modelled as discrete hidden states; the observed data points are realisations of continuous probability distributions with state-specific means that enable ordering of these distributions. The observed sequence is labelled according to the hidden states, permitting only neighbouring states that are also neighbours within the ordering of their associated distributions. The parameters that characterise these state-specific distributions are inferred.

**Results:** We apply our oHMMed algorithms to the proportion of G and C bases (modelled as a mixture of normal distributions) and the number of genes (modelled as a mixture of poisson-gamma distributions) in windows along the human, mouse, and fruit fly genomes. This results in a partitioning of the genomes into regions by statistically distinguishable averages of these features, and in a characterisation of their continuous patterns of variation. In regard to the genomic G and C proportion, this latter result distinguishes oHMMed from segmentation algorithms based in isochore or compositional domain theory. We further use oHMMed to conduct a detailed analysis of variation of chromatin accessibility (ATAC-seq) and epigenetic markers H3K27ac and H3K27me3 (modelled as a mixture of poisson-gamma distributions) along the human chromosome 1 and their correlations.

**Conclusions:** Our algorithms provide a biologically assumption-free approach to characterising genomic landscapes shaped by continuous, autocorrelated patterns of variation. Despite this, the resulting genome segmentation enables extraction of compositionally distinct regions for further downstream analyses.

## 1. Introduction

Hidden Markov models (HMMs) are often described as the workhorse of modern biological sequence analysis [*e.g.,* 13]. While originally used for speech recognition [*e.g.,* 48, 23], they are now central to any field that utilises advanced statistical methods. The first biological application of HMMs was to genome segmentation, in particular to segmentation of genomes according to the level of the bases guanine and cytosine (G+C) vs adenine and thymine (A+T), *i.e.,* segmentation according to GC-rich and GC-poor regions [7, 45].

Generally, HMMs assume an observed sequence that is driven by an unobserved, *i.e.,* a hidden, sequence that in turn is generated by a Markov process. In order to explain the observed sequence, the hidden process and the way in which it generates the sequence of observed data points must be modelled and the parameters of this model inferred. Genome segmentation algorithms in particular infer the marginal probability that each position along a chromosome or stretch of DNA, which corresponds to the observed sequence, belongs to one of a moderately sized number *K* of hidden states, which alternate to form a hidden sequence of states. Traversing the genome on the level of this hidden sequence and considering how to model it, the transition matrix **T***_K×K_* describes the probability of remaining in the same hidden state or switching to another based solely on the most recent state. This dependency on only the most recent previous state makes the model of the hidden sequence a Markov Chain. The sequence of hidden states must be related to the observed sequence, since every data point along the observed sequence is assumed to be emitted conditional on the assigned hidden state at the corresponding place in the hidden sequence. How this relationship is modelled differs between algorithms, but the assignment of each observed genomic region to a hidden state is universally known as “annotation”. In the most classic HMM algorithms, the observed series of data points are assumed to be drawn from a discrete *R* dimensional alphabet according to the matrix of emission probabilities **E***_K×R_* that govern how likely it is for each letter of this alphabet to be emitted by every hidden state. However, they can also be modelled as realisations of a continuous distribution with state-specific parameters such that *e.g.,* the overall distribution of the emitted data conform to a Gaussian mixture model [*e.g.,* 23, 49, 21]. Conditional on both **T** and either **E** or the parameters of the emission densities, the likelihood of the observed data can then be calculated together with the marginal probability of the state at each position using dynamic programming [7, 13], which means that recursive forward and backward passes of an algorithm are performed until parameters that yield a well-fitting model have been inferred. The Baum-Welch algorithm [1, 13], a variant of the expectation-maximisation algorithm, can often be employed to find a local maximum of the likelihood, corresponding estimates 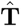, 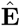, and the initial probability of states 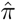. Alternatively, Bayesian approaches typically use Markov Chain Monte Carlo (MCMC) methods, *e.g.,* the Gibbs sampler, to obtain a sample from the joint posterior distribution, from which all marginal posterior distributions follow [*e.g.,* 3, 50, 20, 21].

Our method - oHMMed (ordered HMM with emission densities) – assumes continuous emissions. In one case, the emission is a normal mixture that corresponds to the observed density of the data points. In the other, the emission density is a gamma mixture initially; however, rate parameters of poisson distributions are subsequently drawn from the individual gamma distributions, yielding an observed density of gamma-poisson mixtures (where the data points are discrete counts). Our core assumption for both variants of emissions is that the observed sequence data exhibit appreciable autocorrelation. In order to model this pattern within our HMM framework, the emission densities are parameterised so that they become convex functions within their natural range. This is done by first assuming one shared parameter among the hidden states (the standard deviation for normal distributions, and the shape parameter for gamma distributions), while the other varies between the states (the mean for the normal distributions, and the rate parameter for the gamma distributions). The state-specific parameters can then be used to sort the states by increasing mean of their emitted distributions. Restricting transitions to neighbouring states within the thereby imposed order induces a tridiagonal transition matrix **T** that governs the autocorrelation pattern (and makes the Markov Chain of hidden states reversible). Utilising a Markov Chain Monte Carlo (MCMC) algorithm, oHMMed provides a best-fit annotation of the observed sequence, corresponding estimates of the transition rate matrix, and estimates of the state-specific and shared parameters of the emitted distributions.

The inherent ordering of hidden states by a single parameter bestows oHMMed with several distinguishing properties: Firstly, it avoids the problem of “label switching” that plagues most MCMC methods [32]. Even comparison of output of the same algorithm run multiple times is not straightforward when this occurs; with oHMMed, however, the labels of the states relative to each other are clearly defined and facilitate “label matching” between runs or algorithms. Secondly, the number of estimable parameters is reduced. As expected, we can show that this improves algorithm behaviour and guards against over-fitting. Thirdly, we are able to propose intuitive diagnostic criteria for selecting the appropriate number of hidden states (which is typically assumed as given in classic forward-backward and Baum-Welch algorithms). This is noteworthy because development of model selection criteria for HMMs is complex and no consensus criteria exist [6, 9, 62].

Overall, oHMMed is specifically designed to segment autocorrelated sequences into states that have *statistically significant* differences in mean emissions; these can then be compared meaningfully since they differ only in this metric. This simple and otherwise assumption-free approach is generally applicable whilst remaining agnostic to any causal biological forces; in fact, these can be studied further and without bias after oHMMed segmentation. Previously, general biological autocorrelation patterns have been modelled by incorporating HMM components into more complex econometric and socioeconomic time series models [21]; these describe the observed pattern of stochastic variation (as a “random walk”) with recurring sections where the fluctuations are different means (termed “regime changes”) (see *e.g.,* Markov Switching Models [29] or Changepoint Models, reviewed in [56]).

Recall that the first biological application of HMMs was to the variation of GC proportion along genomes [7, 45]. Even before this, mammalian chromosomes had been described as a “mosaic” of long chromosomal regions of relatively homogeneous GC-content termed “isochores” [12]; these regions are on the order of hundreds of kb to Mb in length. Traditionally, five states of increasing GC proportion within fixed (predetermined) ranges are assumed as part of “isochore theory” [*e.g.,* 8, their Fig 1], and these five states have been delineated in the genomes of various species, including even invertebrates, using specifically formulated binary decision rule segmentation algorithms [5]. Note that the variance in the distribution of GC proportion is also considered to differ between states, *e.g.,* in humans the states of higher average GC proportion are more variable [8]. The validity of “isochore theory” has been fiercely debated [36, 37], especially since “isochores” themselves have been inconsistently defined in terms of their length and level of homogeneity. This prompted the formulation of “compositional domain theory” [16, 15], which posits that the genomic landscape of GC proportion consists of both homogeneous and non-homogeneous regions that can be found by recursive algorithms that maximise the difference in GC proportion between adjacent chromosomal segments using F-tests. In humans, roughly two-thirds of the genome can thereby be classified as consisting of homogeneous segments, but the vast majority are too short to be considered “isochores” [16]. While this dispute around “isochores” has largely been put aside without clear resolution, there is an overall agreement that varying and considerably autocorrelated genomic GC proportions are evident in mammalian sequence data on multiple spatial scales, particularly broader ones: Transitions from regions with high GC content to regions with low GC content generally proceed through a sequence of regions with intermediate GC content *i.e.,* transitions seem to occur predominantly between neighbouring states (see Fig. (1b) in [19] and Fig. (4) in [10]). While the scientific community has not reached a consensus as to the cause of this distinct broad-scale pattern of variation [33, 47], most population geneticists attribute the inhomogeneity of GC proportions *per se* to spatial fluctuation of biased gene conversion [19, 14], which is the preferential use of G and C alleles by DNA repair mechanisms. (Note that even so, the observed *pattern of variation* requires additional explanation.) It has been shown that GC-biased gene conversion can contribute to locally accelerated evolutionary rates [24] and lead to fixation of deleterious alleles [35], thereby impacting genetic inferences and leading to genetic disorders. Thus, being able to identify genomic regions under weaker vs stronger GC-biased gene conversion is still considered important. Our method, specifically oHMMed with normal emission densities, offers an agnostic, probabilistic method to annotate genomes by statistically distinct average levels of GC proportion: It provides a biologically assumption-free characterisation of the continuous pattern of variation, and identifies similar regions that can be extracted for further analysis. We demonstrate this on the genomes of humans, mice, and fruit flies, *e.g.,*using windows of 100 *kb* so as to capture the same observed broad scale variation that led to the past base composition theories (see [53] for an explicit study on the scale of the variation in GC content).

**Figure 1:**
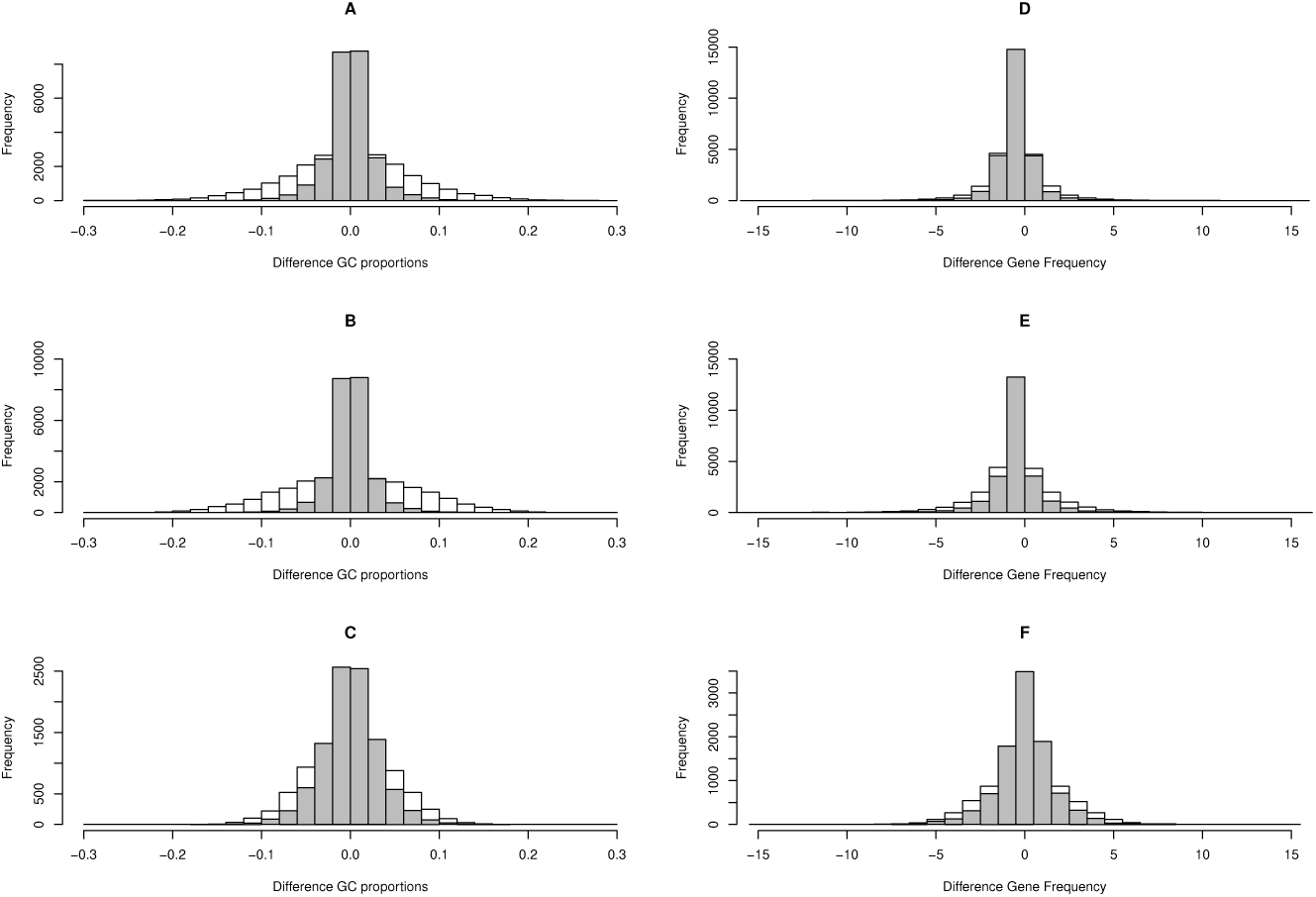
Histograms of pairwise differences in average GC proportion (left column) and average gene content (right column) for neighbouring windows along the observed genomic sequences (grey histograms) and along random permutations of the genomic sequences (white histograms) for humans (A), mice (B), and fruit flies (C) respectively.

Further, we demonstrate usage of oHMMed with gamma-poisson emission densities by annotating the genomes of these same species according to their gene content. Annotations of this kind are useful since information on genomic variation in gene density, or similarly on the density of enhancers regulating gene expression or epigenetic marks may guide inference in studies of biological functions, *e.g.,* regulation of gene expression, cellular differentiation, tissue homeostasis, and response to pathogens. Since gene content is known to correlate with GC proportion [57], we further assess this correlation in our study species using oHMMed output.

Finally, we use oHMMed with gamma-poisson emission densities for a more exploratory investigation of the variation of epigenetic marks along the human chromosome 1; specifically, these marks were sampled from human B cells. We consider three such marks: The first is the output of ATAC-seq (Assay for Transposase-Accessible Chromatin using sequencing) analysis [61].This is a type of assay that identifies and amplifies accessible regions of DNA, aka regions of DNA that are not tightly wrapped around histones (this wrapping results in closed chromatin that cannot be read by the transcription machinery). We will simply call the resulting read counts that make up our data ATAC counts. Importantly, these accessible genomic regions harbour enhancers, insulators, and silencers that determine the finer level of control within the regulatory landscape. The histone modifications deposited as part of the up- and down-regulation of genes are typically subjected to analysis via CHIP-seq (chromatin immunoprecipitation with sequencing) [43, 41], which identifies peak enriched regions of specific histone modifications. Together, more general ATAC-seq analysis and CHIP-seq analysis of precise modification targets can be utilised to assess deviations in the regulatory mechanisms between different cell lines, and how these shift as part of cell differentiation or are altered in disease processes [43, 26].On a broader level, CHIP-seq analyses of core histone modifications within the same cell lines can be combined, *e.g.,* using the multivariate HMM method ChromHMM [18], to segment the genome into differently functional chromatin states, *e.g.,* weak transcription, transcription, poised promoter, flanking promoter. Stacking these analyses across multiple cell lines and issue types produces a comprehensive annotation of the genome [58]. We will consider only two specific histone modifications in human B cells here: H3K27ac and H3K27me3, antagonistic acetylation and methylation marks at the N-terminal position 27 of the H3 histone respectively [44].The mark H3K27me3 is deposited as part of the polycomb repressive complex, an important mammalian gene silencing mechanism [52, 28]. Recruitment and maintenance of polycomb repression is highly context dependent [55], but in the appropriate chromatin environment positive feedback loops can maintain larger repressive domains [30]. There is some evidence that GC-rich DNA is conducive to polycomb recruitment [59]. In detecting H3K27me3 enriched regions, methods must therefore be able to find regions spanning hundreds of *kb* [41]. By contrast, the opposing enhancing mark H3K27ac is know to be enriched in sharp peaks near transcription start sites [41]; however, clusters of peaks known as super-enhancers [31]are highly associated with disease variants. There has recently been evidence that H3K27me3 peaks can cluster similarly [4]. For our purposes, this means that read counts of both marks may exhibit biologically relevant autocorrelated spatial patterns that vary by spatial scale, although they must be negatively correlated with each other across all scales. We therefore decide to analyse the epigenetic marks in 100 *kb* windows (to capture the broader dynamics of ATAC counts and the larger domains formed by both histone modifications) and 1 *kb* windows (to capture the sharper H3K27ac peaks). The results for the 100 *kb* windows can then be related to our previous genomic results to further illuminate the genomic context of the epigenomic landscape.

Overall, this article therefore: (i) introduces oHMMed, a new class of HMMs for autocorrelated sequences, (ii) demonstrates its usage on the well known patterns of genomic base composition and gene content, (iii) and utilises it for a novel study of spatial variation in human epigenetic markers on different spatial scales (100 *kb* and 1 *kb*).

## 2. Materials and Methods

### 2.1. Materials: Genomic Sequences

#### 2.1.1. Human

The *Homo sapiens* GRCh38 reference genome was obtained from the UCSC Genome Browser [34]. The autosomes were subdivided into non-overlapping 100 *kb* windows. For each window with at least 90% successfully sequenced bases (*i.e., >* 90 *kb* non-N bases), the GC proportion was calculated. The number of protein coding genes was determined for the retained windows using GENCODEv21 [22] via the UCSC Genome Browser [34]. Genes were included in a window if the gene coordinates partially or completely overlap with the window coordinates, *i.e.,* genes that overlap into neighbouring windows were not excluded.

#### 2.1.2. Mouse

The *Mus musculus* GRCm39 reference genome was obtained from Ensembl108 [11]. The autosomes were subdivided into non-overlapping 100 *kb* windows. For each window with at least 60% successfully sequenced bases (i.e., *>* 60 *kb* non-N bases), the GC proportion was calculated. Additionally, the number of protein coding genes was counted for each window as before.

#### 2.1.3. Fruit Fly

The reference genome of *Drosophila melanogaster* (version r5.57) was obtained from FlyBase(2022-05) [25]. Since the genome is much shorter than that of the mammals above, the autosomes were subdivided into non-overlapping 10 *kb* windows. For each window comprising more than 90% successfully sequenced bases (*i.e., >* 9 *kb* non-N bases), the GC proportion was calculated. Again, the number of protein coding genes was determined for each window as before.

### 2.2. Materials: Epigenetic Sequences

The sequences of counts of the epigenetic marks ATAC, H3K27me3, and H3K27ac are samples from human B cells. These were obtained from the paired-end alignments (.bam files) on the GRCh38 reference genome provided by ENCODE[17]; the identifiers are ENCSR603LVR for ATAC, ENCSR077YUA for H3K27ac, and ENCSR179LAY for H3K27me3. For each mark, we used only chromosome 1 and aggregated counts into both 100 *kb* and 1 *kb* windows that matched the coordinates of the windows for the previously obtained human genome data.

### 2.3. Methods: Hidden Markov Models with Constrained Transition Probabilities and Emission Densities

#### 2.3.1. General Notation

Assume a vector ***θ*** of random variables, where each entry *θ_l_* represents an unknown or hidden state at position *l* along the genome, where 1 *≤ l ≤ L*. Each *θ_l_* may assume one of *K* states indexed by *i*, with 1 *≤ i ≤ K*. The random vector is a Markovian sequence with an invariant *K × K* transition matrix **T**, where elements Pr(*θ_l_*_+1_ *| θ_l_*) govern the step-by-step probabilities of going from a specific state to the next as the genome is traversed. Depending on the state probabilities at every position along the genome 1 *≤ l ≤ L*, a realisation (data point) *y_l_* is sampled from a (continuous) probability density function. These are collected in the vector of realised emissions ***y***.

#### 2.3.2. Convex Emission Densities

They key assumption behind oHMMed is that the hidden states can be ordered in a way that is reflected in their emissions. To illustrate: assume two states *θ_i_*, *θ_j_* with (*i, j*) *ɛ* [1*, .., K*] and *i < j*, and two emissions (*y_i_, y_j_*) *ɛ* ***y*** with *y_i_ < y_j_*. Then the relationship:

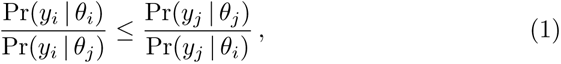

should hold, and hold with equality only on a null set. In other words, the ratio 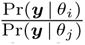 should be convex. For strictly positive densities, we can take the logarithm here and define a convexity condition that must be fulfilled for an ordering of states to be possible:

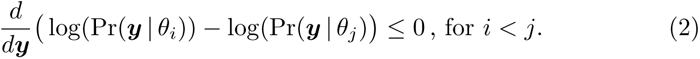

##### Normal Emissions

Consider any two of *K* total states that each emit normal distributions; in particular *y_i_* is drawn from *N* (*µ_i_, σ*^2^) and *y_j_* is drawn from *N* (*µ_j_, σ*^2^) where *µ_i_ ≤ µ_j_*. The above convexity condition holds:

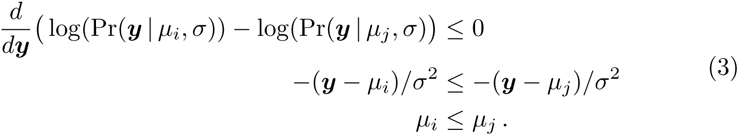

Note that the shared standard deviation *σ* between states is a necessary assumption here.

##### Gamma Emissions

Consider any two of *K* total states that each emit a gamma distribution; in particular *y_i_* is drawn from *G*(*α, β_i_*) and *y_j_* is drawn from *G*(*α, β_j_*) where *β_i_ ≤ β_j_*. The above convexity condition holds:

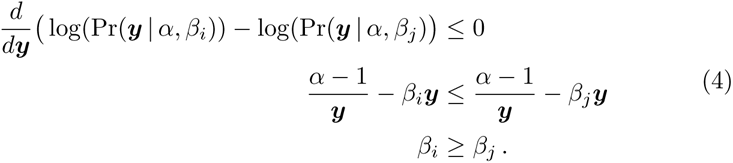

Here, the shared shape parameter *α* between states is a necessary assumption.

#### 2.3.3. Transition Probabilities

In addition to assuming that the hidden states can be ordered, we restrict transitions to neighbouring states within this ordering. This results in a tridiagonal *KxK* transition matrix **T**, since Pr(*θ_i_ | θ_j_*) *>* 0 only for *j* in (*i−* 1*, i, i* + 1), while Pr(*θ_i_ | θ_j_*) = 0 otherwise. Note that the transition matrix thus has 2*K −* 2 estimable parameters. Further, we let the prior probabilities of the hidden states correspond to the stationary distribution of the transition matrix **T**, which we denote as the row vector ***π*** with entries *π_i_* = Pr(*θ_l−_*_1_ = *i*). It follows that the system is in detailed balance, *i.e.,* fulfills the equations:

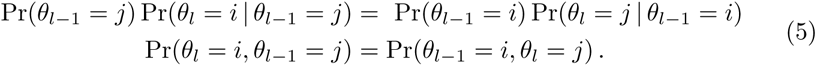

Note that this corresponds to the structure of double-stranded DNA sequences, where the 5’ end of one strand corresponds to the 3’ end of the other.

#### 2.3.4. MCMC Algorithm

The assignment of each position along the genome to a hidden state is determined by the forward-backward passes of a HMM algorithm (see Additional File 1, Section “Forward backward HMM algorithm“). After each pass, the transition rates and parameters of the emission densities per state must be estimated and their fit evaluated. Baum-Welch expectation-maximisation algorithms, which are often employed for parameter estimation with HMMs [*e.g.,* 13], require independent maximisation of ***π*** and **T**. Therefore, we develop a Markov Chain Monte Carlo algorithm, in particular a Gibbs sampler, that estimates the posterior distributions of the transition rates and parameters of the emitted distributions given the current annotation and the observed data. The samplers are fully characterised in Additional File 1, Section A1 for normal and Additional File 1, Section A2 for gamma-poisson emissions. We also provide a graphical description of the different versions of the oHMMed algorithms in the respective files, which may facilitate understanding of the algorithm structure.

#### 2.3.5. Implementation

Our algorithms are available as the R package oHMMed on CRAN [38], with the source files also deposited on GitHub [39]. Explicit usage recommendations [40] can also be found as a manual on GitHub, which include pointers on setting partially informative priors and initial values for the estimable parameters. We also describe the accompanying suite of diagnostics for assessing convergence and model fit. Note that oHMMed performs 10 iterations of the Gibbs sampler in roughly 1.06 seconds on a sequence of length 2^11^, and that the speed decreases linearly with sequence length (details in the usage recommendations on GitHub[40]).

## 3. Results and Discussion

### 3.1. Empirical Transition Rates

At the outset, it must be ensured that data conform to oHMMed assumptions. We illustrate in depth how to compare the mean differences in average GC proportion and in protein coding gene content of consecutive windows along the genomes of our study species to the same differences in random permutations of these windows to check for autocorrelation (see Fig. 1). The variance of differences for the genomic GC proportion in humans, mice, and fruit flies respectively is 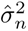 = (0.000673,0.00055,0.00103) compared to 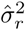 = (0.00469, 0.00552, 0.0021) for the permuted genome sequence, with the ratios being highly significant in all cases (F-test with p-value *p <* 2.2*e^−^*^16^). Similarly, we report variance differences of 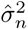 = (1.14,1.73,2.54) in the number of protein coding genes in consecutive genomic windows of the three species compared to 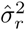 = (2.43,5.01,4.29) in the random sequence, attaining the same level of significance for these ratios. Overall, we find evidence for an underlying autocorrelation structure in the observed sequences of both base composition and gene content in all species. Note, however, that this pattern is particularly pronounced in the genomic GC proportion of humans and mice (see Fig. 1A, B). We perform similar checks on the sequences of epigenetic marks, and find that the difference in variance between counts in consecutive windows is considerably lower than expected for randomly arranged windows (F-test with p-values always below *p <* 9.7*e^−^*^21^) However, we omit the in depth reporting here so as not to be overly repetitive.

### 3.2. Segmentation of the Human Genome

#### GC Proportion

After running oHMMed with normal emission densities on the human genomic GC proportion in 100 *kb* windows several times assuming *K* = (2*, ..,* 8) hidden states, we chose to segment the GC content into *K* = 5 states (all runs with 1500 iterations and a 20% burn-in). This is a compromise between two aspects of the inference procedure that we have chosen as diagnostics: The first is model fit. Adding more states one by one starting from *K* = 2 will - at least initially - lead to an increasing posterior log-likelihood as the overall emitted distribution becomes more flexible. However, there should be a number of states after which the increase in log-likelihood plateaus, pointing to a candidate number of hidden states. The second aspect of the decision concerns whether the difference between means of the emissions generated by neighbouring states are statistically different. For the human GC proportion, the posterior log-likelihoods start to plateau after about *K* = 5 (Fig. 2A) and the means are well separated, *i.e.,* the 68% confidence intervals (means plus/minus one standard deviation) barely overlap (see Fig. 2C). In fact, only first and second states clearly overlap at this level, and the second and third do so marginally; however, this interpretation of the plots is conservative and the means are still statistically distinguishable by a standard (one-sided) t-test at the 95% confidence level (part of standard oHMMed output). From Fig. 2C, the continuous variation of observed GC content among windows is also evident: Discretisation into *K* = 5 states introduces discontinuities, in particular between the wider-spaced means. Increasing *K* would have resulted in an increasing overall likelihood (Fig. 2A) and better fitting overall distribution of emissions compared to the distribution of observed GC proportion along the genome (see Figs. 2C, D) at the cost of increasing the overlap between neighbouring states to the point of non-significant t-test outcomes. Decreasing *K* would have the opposite result.

**Figure 2:**
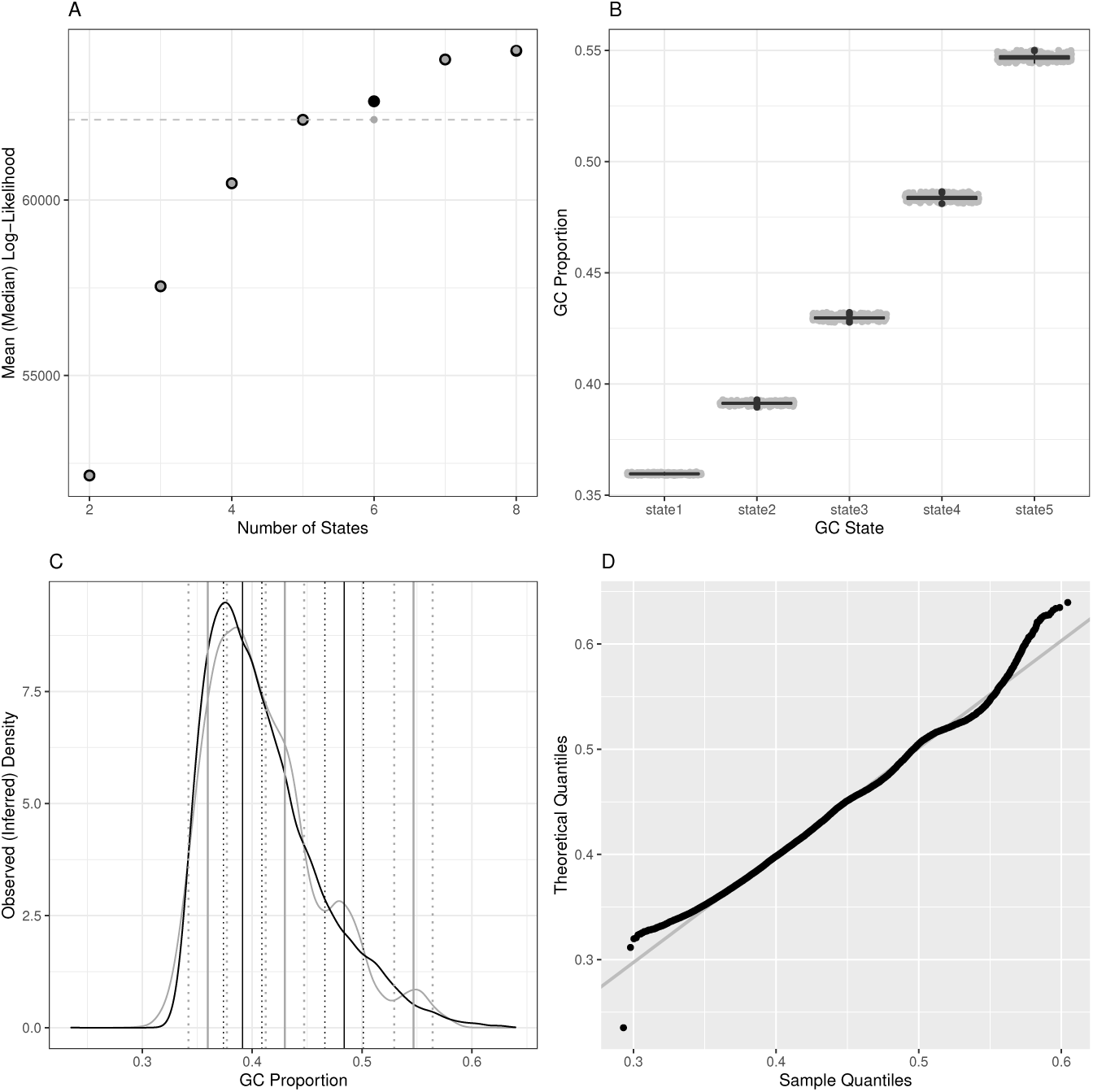
Summarised diagnostics for annotation of the human genome by average GC pro-portion using oHMMed with normal emission densities. Panel A shows the mean (black) and median (grey) log-likelihood of fully converged runs of the algorithm with different numbers of hidden states, with the dashed horizontal line marking the selected number of hidden states, which is five. The difference between the mean and median for six hidden states is the effect of autocorrelation in the traces of the estimated parameters. In panel B, boxplots of the posterior (*i.e.,* inferred) mean GC proportion of the run with the five hidden states are presented. Panel C shows the observed overall density (black) of the GC proportion superimposed on the posterior (inferred) density, with the inferred means per chosen number of states plus the 68% confidence intervals drawn in vertical lines. The final panel D shows the QQ-plot of the observed density vs. the posterior density (here termed the theoretical distribution). Full descriptions of the diagnostics available for oHMMed can be found in our usage recommendations on GitHub[40], and the code for this visual summary is available as an R script named “oHMMedOutputAnalyses.R”[39] on GitHub.

For the selected five states, we inferred means of 0.360,0.391,0.430,0.4834,0.567 respectively, and a shared standard deviation of 0.174. The proportion of genomic windows assigned to each state splits to 0.261, 0.320, 0.256, 0.125, 0.038. On average, 29, 9, 5, 3 and 4 successive 100*kb* windows fall within the same hidden state for increasing GC proportions respectively.

Generally, segmentation of the human genome into 5 states at this scaling aligns with past literature [8, 10], with ”isochore theory” dictating a split of comparatively homogeneous DNA regions *≥* 300*kb* into classes with predefined mean GC proportions of *<* 0.38, 0.38 *−* 0.42, 0.42 *−* 0.47, 0.47 *−* 0.52*, >* 0.52 based on both the variation of mean and standard deviation of G+C alleles. Note that each of these categories contains one of our inferred means. Relative proportions of human DNA in these classes are cited as 0.19, 0.37, 0.31, 0.11, 0.03, with the lowest two classes often merged; this differs somewhat from our inference.

Our method additionally infers a transition rate matrix of

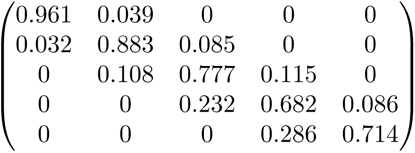

between the hidden states. This, together with our inferred means, reflects the often referenced mosaic structure of hominid genome sequences: Broad troughs of regions with low GC content that transition into regions with means that are almost comparable, and narrow rugged peaks of regions with high GC with more frequent transitions into neighbouring states with increasingly differentiated means (Fig. 2B, Fig. 6A). An exception to this landscape is the left arm of chromosome 1, which exhibits a wide region of high GC content (see Fig. 6A).

#### Gene Content

We used oHMMed with gamma-poisson emission densities to segment the human genome according to the number of protein coding genes per consecutive window several times assuming *K* = (2*, ..,* 5) hidden states (all runs with 40000 iterations and a 12.5% burn-in). Our diagnostics suggest *K* = 3 hidden states, since this is where the increase in log-likelihood compared between the runs begins to taper off and discrimination between the state-specific means is statistically possible (see Fig. 3). A reasonable fit to the observed histogram of overall counts is achieved by the (smoothed) theoretical curve inferred by this model (see Fig. 4); there is some overestimation of the occurrence of zero counts and underestimation of the single counts.

**Figure 3:**
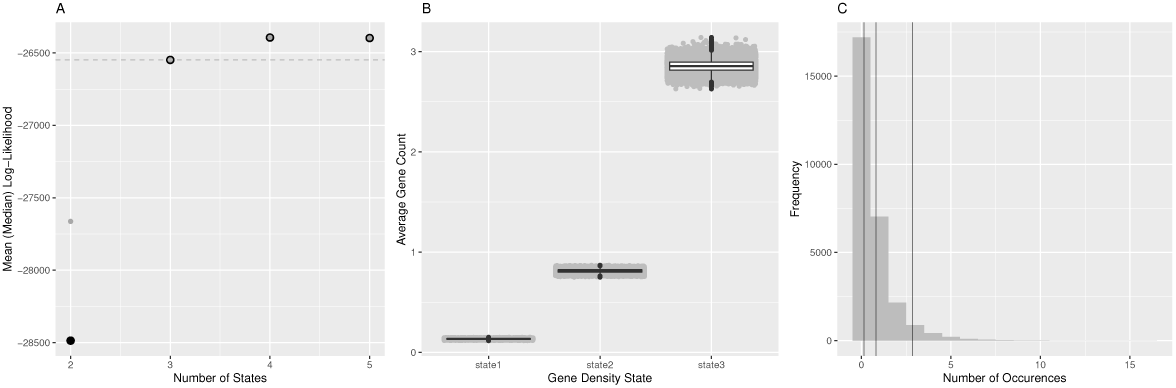
Here we show the first part of the summarised diagnostics for oHHMed with gamma-poisson emission densities as employed on counts of the average number of protein coding genes along the human genome. Panel A shows the mean (black) and median (grey) log-likelihood of fully converged runs of the algorithm with different numbers of hidden states, with the dashed horizontal line marking the chosen number - which is three. In panel B, boxplots of the posterior (i.e. inferred) mean gene densities of the inference run with three hidden states are presented. Panel C shows the observed distribution of gene counts with the inferred means superimposed as vertical lines. These are significantly different on the 95% confidence level as per one-sided poisson rate test (part of standard oHMMed output). Once again, full descriptions of the diagnostics available for oHMMed can be found in our in our usage recommendations on GitHub[40], and the code for this visual summary plus the corresponding rootogram is available as an R script named “oHMMedOutputAnalyses.R”[39] on GitHub.

**Figure 4:**
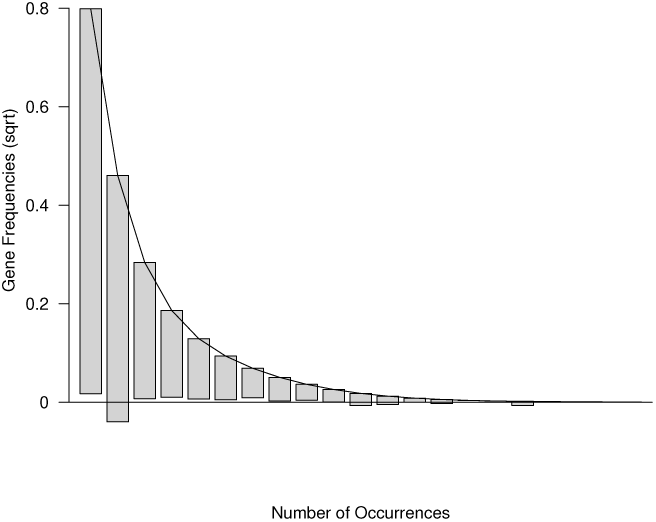
As the final part of the summarised diagnostics for oHMMed with gamma-poisson emission densities and three hidden states as applied to the protein coding genes in humans, we present the above rootogram: The bars represent the observed frequency of counts (square root transformed), and they have been shifted so that the top of each bar aligns with the (smoothed) distribution inferred by oHMMed. Deviations can therefore be assessed by checking the distance of the lower end of each bar to the x-axis.

The landscape of protein coding genes is not as varied as that of the GC proportion: Longer regions of both low and high gene counts occur with a slight abundance of the former; all show similar propensity to transitioning to neighbouring states (see Fig. 6B), as is further evidenced by the inferred transition rate matrix:

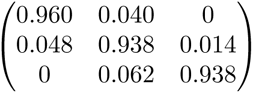

Note that this translates to an average of 31, 20 and 19 subsequent windows being assigned the same state in order of increasing gene content. The inferred means of 0.094, 0.751, 2.80 per state with the proportions of windows assigned to them being 0.499, 0.412, 0.089 respectively indicate that genes are generally sparse with mostly only one or none at all per window.

#### Correlation

Although the GC proportion and the density of protein coding genes appear to vary on a different scale, counting the number of occurrences of each of the 5 GC states within the 3 states for gene density reveals a clear positive correlation (see Fig. 5A), and compare [5, Fig. 7]. In fact, the position-specific posterior means of the GC content and the gene density, *i.e.,* taking the sum over the inferred means times the probabilities of being assigned to the respective state at each position along the genome, have a highly significant (p*<* 2.2*e^−^*^16^) positive Spearman correlation coefficient of 0.608. Running versions of this correlation can be applied to screen for anomalous regions (see Fig. 6C, and note the negative correlations around most telomeres).

**Figure 5:**
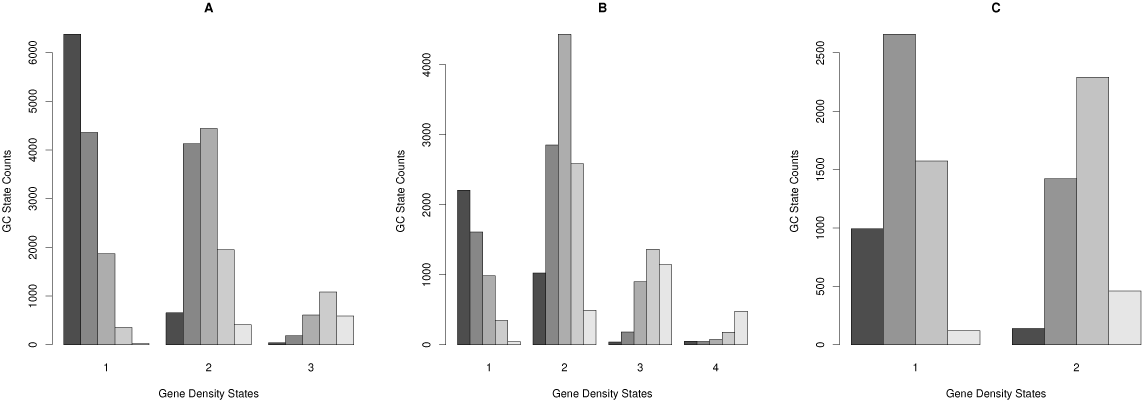
Barplots that visualise the cross-tabulation of the two genome annotation results per species: For the full genome of the human (A), mouse (B), and fruit fly (C) respectively, we count the number of windows assigned to each GC state (y-axis) within regions assigned to each gene density state (x-axis) and show them in decreasing shades of grey. Essentially, this is a discretised representation of the positive correlation between these features.

**Figure 6:**
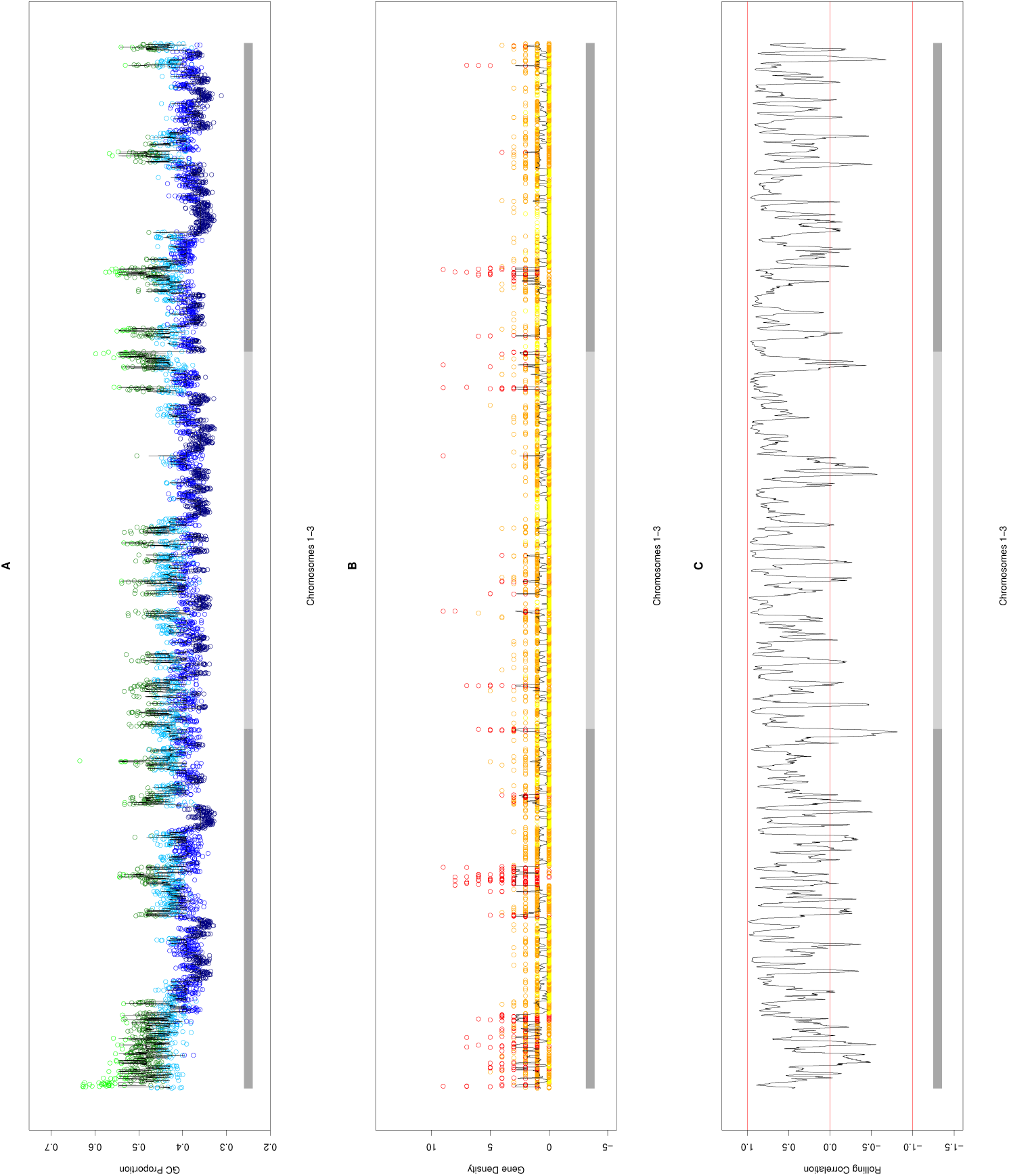
In each of the above panels A-C, the human chromosomes 1 *−* 3 (demarked by alternating dark and light grey horizontal bars) are plotted for different oHMMed analyses: In A, the average GC proportion in every 100 *kb*window is coloured by the oHMMed-inferred GC state. The three lower states are in blue and the two higher ones in green, with the shades lightening with higher GC proportion. In B, the number of protein coding genes for every 100 *kb* window are shown in colours corresponding to the oHMMed-inferred gene density states: yellow, orange, and red mark increasing gene density states. Note that in both A and B, the black lines trace the posterior (inferred) means returned by oHMMed with normal and gamma-poisson emissions respectively. These position-specific posterior means are the sum of estimated means times the respective probabilities of each state, thus combining both estimated mean values and the algorithm’s certainty of the assigned state. In C, Spearman’s correlation for the two posterior means in shown in rolling windows of 40 collated 100 *kb* windows.

**Figure 7:**
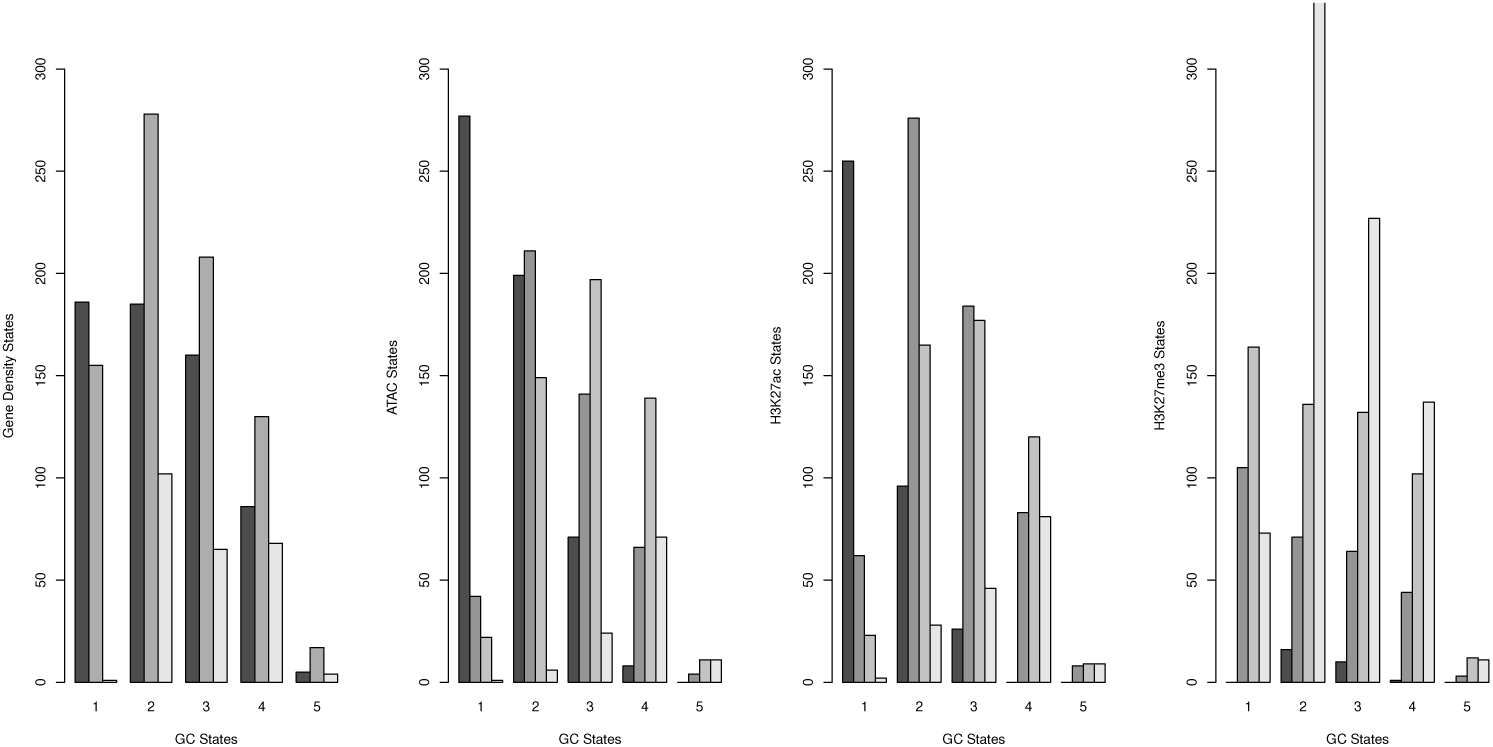
The above barplots visualise the cross-tabulation of the annotation of the human chromosome 1 by GC content (x-axis) with the annotation by gene density, ATAC, H3K27ac, and H3K27me3 respectively (y-axes). More specifically, we count the number of 100 *kb* windows assigned to each state per feature on the y-axes within regions assigned to each GC content state (x-axis), and show them in decreasing shades of grey. Essentially, this is a discretised representation of the correlation between the GC content and the respective other genomic and epigenomic features.

### 3.3. Segmentation of Mouse and Fly Genomes

The genomic landscape of the mouse is generally more homogeneous and compact than that of humans, with comparatively narrower troughs and less rugged peaks in the variation of GC proportion and a wider range in gene counts per window. Segmentation results in regular, distinct genomic regions (*K* = 5 for the GC proportion and *K* = 4 for the gene content, with the same number of iterations and burn-in percentage as previously; see Additional File 2 Section “Segmentation of the Mouse Genome“), and a cleaner discretised correlation pattern (see Fig. 5B).

In the much more condensed genome of the fruit fly, clean segmentation based on smaller genomic windows is achieved with fewer hidden states; the results are primarily indicative of chromosomal structure. Specifically, we infer *K* = 4 for the GC proportion and *K* = 2 for the gene content (again with the same number of iterations and burn-in percentage; but see Additional File 2 Section “Segmentation of the Fruit Fly Genome“ for details). Despite comparatively less quantifiable variation in either feature, positive correlation between them is still apparent (see Fig. 5C).

### 3.4. Segmentation of Human Chr1 By Epigenetic Marks

We applied oHMMed with gamma-poisson emission densities to the epigenetic marks along human chromosome 1, with counts parsed into both 100kb and 1kb windows. Importantly, we decided to remove telomere- and centromere-adjacent regions from our analyses since these genomic regions have their own unique dynamics. For the 100 kb data, this amounted to 200 removed windows from the chromosome ends and 204 additionally removed windows from the left and 203 from the right side of the centromere (process performed by visual assessment of the distribution of the counts of the remaining windows for outliers); the same number of windows times 100 were removed for the 1kb data. The procedure of running the oHMMed algorithm on the resulting sequences was the same as described in the previous subsections for the genomic data. Therefore, we will focus on the biological outcomes in the main text and show the core results pertaining to the running of the algorithms in the Additional File 4.

#### 3.4.1. Results for the Broad Scale 100 kb Windows

oHMMed analysis indicated that *n* = 4 hidden states are appropriate for the ATAC, H3K27ac, and the H3K27me3 counts (see Table 1 in Additional File 4). The epigenetic landscape of ATAC and H3K27ac counts exhibit a pattern familiar to us from the genomic landscapes (see Tables 2,3 as well as Figure 1 in Additional File 4): Long regions of less accessible chromatin, with fewer epigenetic modifications, are punctuated by shorter peaks corresponding to more accessible chromatin/enrichment of modification marks. In the case of H3K27ac, the states with increasing counts are also increasingly transient. The landscape of H3K27me3, however, is fascinatingly different (see Table 4 as well as Figure 1 in Additional File 4): It consists to a large part of the most highly enriched state, which is inherently variable but forms a sort of hilly plateau. This is interrupted by the less highly enriched regions, whose length decreases with enrichment level. Notably, the state with the lowest average number of counts is comparatively devoid of signal compared with the lowest state of the other marks. Note that these results for H3K27me3 are actually pleasantly inline with the fact that it is known to have broad enrichment peaks, as described in the Introduction (Section 1); by comparison, H3K27ac varies primarily on a shorter scale and amongst lower enrichment levels and, while the windows in the highest enrichment state may harbour clusters of known super-enhancers, it is beyond our scope to investigate this further here.

Comparing epigenetic marks amongst each other and within the genomic context, we find the expected negative correlation between the antagonistic marks H3K27ac and H3K27me3, which is borderline statistically significant, as well as equally biologically plausible, strongly significant correlations between ATAC, H3K27ac, and gene density (since only transcriptionally accessible genomic regions can harbour active genes); see Figure 8A. These correlations further imply a negative correlation between H3K27me3 and ATAC as well as between H3K27me3 and gene density. In analysing the correlations separately within every oHMMed-inferred GC state in Figures 8B-F and Figure 7,we find the following pattern: Regions of low GC content, which are largely devoid of protein coding genes, contain higher levels of the silencing H3K27me3 than any other epigenetic mark. These regions appear generally inactive, and all marks are positively correlated. Then, as the GC content increases across states, so do the histone modifications and the gene density, and the previously described correlations appear and become more statistically significant with increasing GC content. In the highest GC state, there appears to be very little data and the correlations thus become weaker and less significant. Thus, we validate the importance of the genomic context in analysing histone modifications.

**Figure 8:**
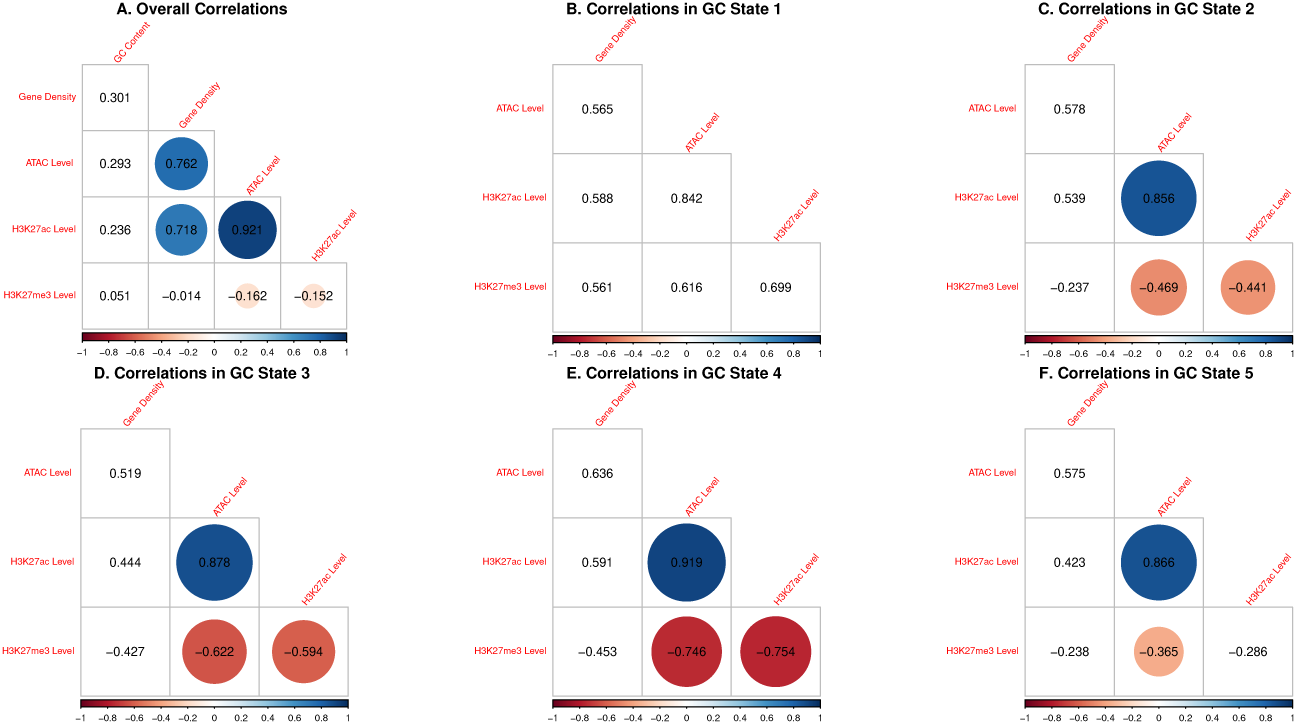
The above figures show Spearman’s correlation between the posterior means (which are the sum of the estimated means times the respective state probabilities) between all analysed genomic and epigenomic features on human chromosome 1 in 100 *kb* windows; correlations that are significant on a 0.99 significance are circled. In panel A, the overall correlations are shown. In panels B-F, correlations between all remaining features are shown separately for regions of different GC content (from low to high).

#### 3.4.2. Results for the Fine Scale 1 kb Windows

oHMMed analysis here indicated that *n* = 5 hidden states are appropriate for the ATAC counts, and *n* = 7 were suitable for the H3K27ac and the H3K27me3 counts (see Table 1 in Additional File 4). Straight away, we would like to note that the segmentation according to 1 *kb* windows picks up very different signals to segmentation according to 100 *kb* windows, and it is not apparent how to easily relate the two spatial scales. Essentially, the 1 *kb* segmentation tracks many more slight changes in the landscape, describing the regions of low epigenetic activity in greater detail. The finer epigenetic landscape of ATAC counts is the most variable amongst the three marks; the third ATAC state, which corresponds to slightly accessible chromatin (judging by the inferred mean), forms the only true stable ATAC domain (see Table 2 as well as Figure 2 in Additional File 4). For H3K27ac, the oHMMed algorithm partitions out regions practically devoid of enrichment for the lowest state; the states corresponding to regions with low enrichment levels are the most stable, and states of increasing enrichment then becoming increasingly more variable (see Table 3 as well as Figure 2 in Additional File 4). The finer epigenetic landscape of H3K27me3 is defined by comparatively longer stretches of more the same state than are present for the other marks (see Table 4 as well as Figure 2 in Additional File 4). The first inferred state for H3K27me3 also corresponds to regions in which histone modifications are essentially absent, and it forms occasional troughs in the landscape. However, every level of enrichment of H3K27me3 is well-represented by decently stable regions in the genome, particularly the state corresponding to the second highest enrichment level. Overall, we therefore again see that H3K27me3 varies in a more modulated manner across larger spatial scales than the other marks.

The finer segmentation of epigenetic marks lends itself to interpretation within the context of functional genome annotation; we will distinguish between windows that fall solely into intergenic regions, gene bodies, promoters, as well as gene bodies and promoters, since these categories were given in the data files from which we obtained the marks themselves) (see Table 9), Note that by comparison, 100 *kb* windows will typically never contain just a promoter. Overall, there is once again a high positive correlation between ATAC and H3K27ac and a lower negative correlation between these and H3K27me3; however, these correlations are not deemed significant, perhaps because the fine scaling introduces too much variability (see Figure 10A.) When assessing the regions of different functional annotation separately, the strongest correlations are evident in regions that contain promoters, with a strong and significant negative correlation between H3K27ac and H3K27me3, and a high positive correlation between ATAC and the former (see Figure 10C,E). This appears biologically intuitive, since histone modifications should be located to regions proximal to promoters and these must further be read by the transcription machinery in order to influence gene expression. Weaker but otherwise similar correlations can be observed in windows that fall within genes (see Figure 10D).

**Figure 9:**
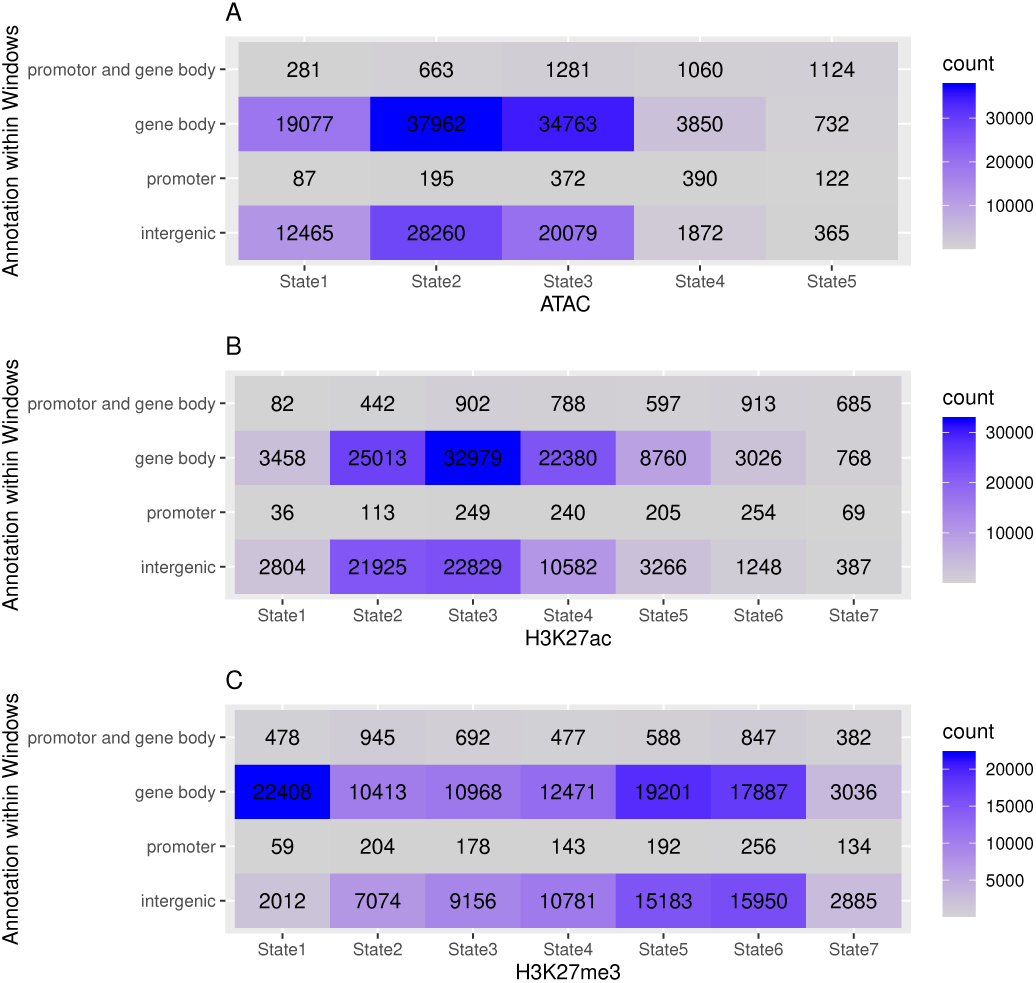
In lieu of barplots (which are sub-optimal in this case because of the extreme differences in counts between cells), we here present tables that show the number of 1 *kb* windows per oHHMed-inferred epigenetic marker state (columns) that fall within specific functionally annotated regions (rows). Panel A shows results for ATAC, panel B those for H3K27ac, and panel C those for H3K27me3; in all panels, the cells in the tables are coloured by darkening shades of purple for increasing counts.

**Figure 10:**
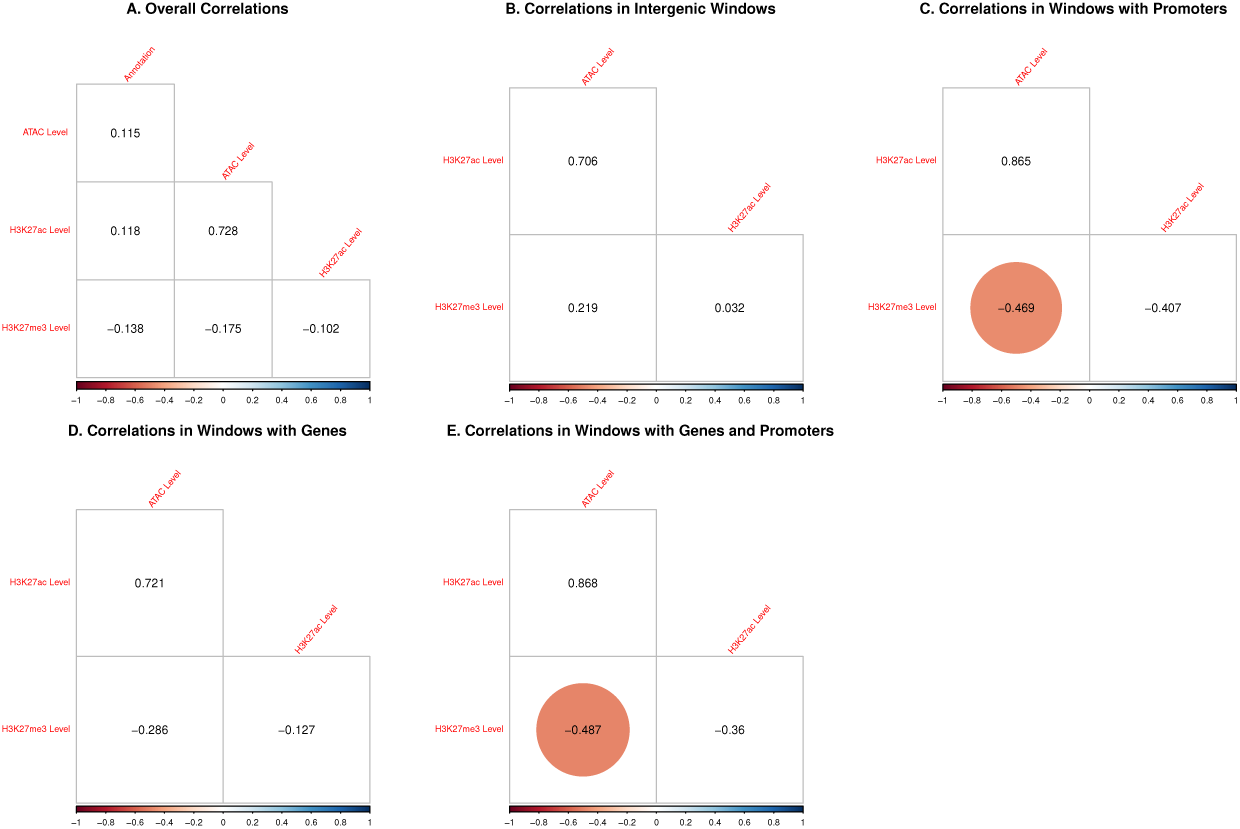
The above figures show Spearman’s correlation between the posterior means (which are the sum of the estimated means times the respective state probabilities) among the epigenetic marks as well as the functional annotation on human chromosome 1 in 1 *kb* windows; correlations that are significant on a 0.99 significance level are circled. In panel A, the overall correlations are shown. In panels B-F, correlations between the marks are shown separately for regions with different functional annotations.

### 3.5. Comparison to HMM with Unordered States

Recall that, in contrast to HMM algorithms with unordered states, the development of oHMMed has: (i) reduced the variability of state-specific emitted densities from 2*K* to *K* + 1 parameters, (ii) restricted the transitions between hidden states so that the transition rate matrix is tridiagonal resulting in a reduction from *K*^2^ *− K* to 2*K −* 2 parameters, and (iii) used reversibility to determine the prior distributions of states instead of specifying *K −* 1 parameters. Thus, rather than the full *K*(*K* + 2) *−* 1 estimable parameters, we are left with 3*K −* 1.

Despite greater flexibility, the unordered versions of the oHMMed algorithms perform either no better than oHMMed (in the case of normal emissions) or worse than oHMMed (in the case of poisson-gamma emissions) in terms of the overall posterior log-likelihood and percentage of correctly assigned states when the algorithms are pitted against each other on simulated sequences with inherent autocorrelation (see Additional File 3 Section “Results: Summary and Interpretation“). While no model selection criterion for comparing HMMs with fixed numbers of states but different numbers of parameters exist, it can be deduced that classic model selection criteria that penalise overall model fit determined via maximum-likelihood by model complexity as determined via the (effective) number of parameters (such as AIC, BIC, and DIC) would likely favour oHMMed to guard against over-fitting.

More importantly, oHMMed is designed to prioritise segmentation into regions with statistically different mean emissions. Indeed, it finds such partitions more readily than its unordered counterpart, particularly on shorter sequences than typically used in full genome analysis (again, see simulations in Additional File 3 Section “Results: Summary and Interpretation“). For the human genomic sequences analysed in this article, which are very long, both methods yield similar estimates (see Additional File 3 Section “Results: Hominid Data“).

It is also important to test oHMMed on sequences that clearly violate its model assumptions, for example on sequences simulated using its unordered counterpart (see Additional File 4). If hidden states have very different standard deviations, oHMMed incurs a predictable bias in estimation of the state-specific means since it can infer only one shared standard deviation and this is typically near the average of the true standard deviations. If the inferred standard deviation is comparatively high, oHMMed may effectively merge hidden states and infer fewer effective states than truly present in the data. Overall, its unordered counterpart therefore consistently infers an overall better-fitting model as determined by average posterior log-likelihood. (Recall again that there is no suitable metric for comparison of these HMMs, but the difference in average posterior log-likelihood is great enough that more refined measures should hardly change the outcome). However, it is worth mentioning that deviant behaviour by oHMMed in the cases we tested is no more frequent than mis-inference by its unordered counterpart due to the larger number of estimable parameters. The latter clearly requires very large data sets, high numbers of iterations, and full diagnostics to perform well, even in a controlled setting.

Overall, we therefore recommend testing for autocorrelation as in Section 3 before application of oHMMed, and only to do so if it is present. If, in post-analysis, our recommended diagnostics indicate that hidden states have very different standard deviations (the observed vs inferred density plots may show this, or the prevalence of fewer effective inferred hidden states than set in the algorithm), one should consider using unordered algorithms. However, if this is not the case, oHMMed is a robust and accurate algorithm.

## 4. Conclusions

In this article, we developed algorithms for characterising the large scale variation in genomic data. Part of the inspiration was provided by the clear visual indication that the genomic GC proportion of hominids likens a continuous, reversible random walk with a finite number of re-occurring changes in mean. Furthermore, when partitioning the genome into windows, anecdotal evidence abounds that [10]: “very large GC differences at borders [… are] rare, thus leading to the formation of blocks […] from closer [… GC levels]”. The continuous nature of this observed pattern contrasts with some formulations of the long-standing “isochore theory”, which postulates homogeneous stretches of (five, or sometimes four) discrete states (aka “isochores”) with sharp boundaries between them. Based on “isochore theory”, the program IsoFinder [42] and related methods [51] use a binary decision rule to sequentially “slice” genomes into piece-wise constant sections of different means; they have been heavily criticised for finding “isochores” that are not actually there [27]. A closer fit to the observed continuous pattern of variation is provided by Fearnhead and Vasileiou [21], whose Bayesian Online Changepoint model simultaneously infers all the points within the observed sequence at which the underlying assignment of hidden states changes from one to another through direct simulation. The algorithm infers a smoothed mean GC content across the observed sequence by averaging over the likelihood of assignment to ”isochore states”, and does so in a run-time that scales quadratically with sequence length.

Differently to all the above, the central aspect of oHMMed is definitive sequence segmentation or annotation that is agnostic to the number of hidden states, as well as to the causal forces of the observed sequence pattern. Despite using piece-wise constant approximations to the data purely for methodological reasons, we are able to model autocorrelation patterns by ordering hidden states according to the means of their emission densities and restricting transitions to neighbouring states. By doing so, oHMMed specifically captures the pattern of stochastic variation with underlying regime changes observed in genomic sequences of GC proportions that is not specifically represented in “isochore theory” and provides a descriptive quantification of these patterns, although it also recovers 5 states of statistically similar average GC content. However, the assumptions made to model autocorrelation, which in itself appears to be an empirically well-founded observation, necessitate equal variances between state-specific emitted distributions, which is generally not given exactly. We have shown that, if variances are quite different, oHMMed will either infer state-specific means with a predictable bias or merge states and fit the corresponding model. However, deviations in variance between states that are not extreme will still lead to sufficiently accurate results.

Note that the sequence data required as input for oHMMed algorithms must already be partitioned into windows, which again distinguishes it from the other algorithms. We believe that the window size should be chosen according to the biological research question: For the sequences of genomic data in this article, the window sizes were set to best illustrate the considerable patterned variation in GC proportion and gene density along the genomes of the study species compared to their length. Specifically, this means that the 100 *kb* spatial scale analysed is comparable to that of the length of “isochores”, although “compositional domain theory” argues that the majority of regions that can be classed as having homogeneous GC proportions may be shorter than our genomic windows [16]. We posit that specifically altering the size of the genomic windows in the input data could form the basis of comparative studies of the genomic variation of GC proportion across different spatial scales, since the causes and implications of variation may differ between these [53] and have to date not been fully uncovered. In order to do this, it is crucial to be able to extract distinct genomic regions for subsequent analyses without getting embroiled in old debates. We propose oHMMed as a powerful, assumption-free tool for this.

The window sizes for the epigenetic data were selected in part to illustrate how altering these can uncover different spatial dynamics and be incorporated into comparative studies involving other genomic features: On the 100 *kb* scale, which is known to be appropriate for epigenetic marks with broad enrichment peaks such as H3K27me3, we are also able to conjointly assess variation in the epigenetic landscapes and the genomic landscapes of GC content and gene density. The 1 *kb* scale is appropriate for the sharper peaks in the profiles of epigenetic marks, and enables a joint assessment with functional genome annotations.

From an overall modelling perspective, oHMMed falls within the traditional HMM framework familiar to most bioinformaticians. The accompanying suite of diagnostics, particularly for finding the appropriate number of hidden states, is straightforward and intuitive. In fact, the lack of complexity makes oHMMed preferable over even a standard unordered HMM with the same underlying MCMC sampler in terms of model fit, particularly when the emission densities are not normally distributed. Since its run-time scales linearly with sequence length, application to long sequences is feasible.

Beyond oHMMed’s appeal in methodological tractability and descriptive analyses, we would like to emphasise the interpretability of its output: Since hidden states can be definitively compared by their mean emission densities (as it is the only metric they differ in), segmentation of sequences into regions with statistically significant average patterns of variation is possible. Recall that these states can therefore also be “label matched” between runs, both on the same and on different data sets.

We would like to stress that oHMMed is a generalised method, which distinguishes it from often more complex and fine-tuned methods developed for specific genomic features (*e.g.,* the GC proportion [42] or the recombination rate [60]). In this article, we developed oHMMed with normal emission densities and applied it to the window-based genomic GC proportion of humans, mice, and fruit flies; however, it could also be employed for, *e.g.,* sequences of average recombination rates per genomic window (after normalising transforms), or window-based measures of epigenetic marker counts with such broad enrichment peaks (spanning *Mb*s) that their distributions approach normality (pending, of course, checks for autocorrelation in these features). Further, we extended oHMMed to gamma-poisson emission densities, which we first applied to the protein-coding genes of humans, mice, and fruit flies. We then utilised this version of oHMMed to analyse the patterns of variation in the epigenetic data given by ATAC-seq read counts and the CHIP-seq read counts of the markers H3K27ac and H3K27me3. It could likely be run not only on the many other epigenetic markers and transcription factor binding sites that can obtained via CHIP-seq analysis, as well as sequences of window-based count data pertaining to other regulatory genomic features such as the number of promoters, enhancers, or repressors.

Since oHMMed makes no biological assumptions and the inferred hidden states can be ordered, it also facilitates analyses of associations between genome segmentations performed according to different features. In this article, we initially simply show the positive correlation between genome annotations by GC proportion and protein-coding gene content in 100 *kb* windows. This is done by simply cross-tabulating the assignments of genomic windows to hidden states, as well as by correlating the position-specific posterior means; running versions of the latter can be applied to screen for regions of interest. More importantly, the fact that we used the same algorithm and the same window sizes on the epigenetic data enabled us to assess the correlations between ATAC counts, H3K27ac, and H3K27me3 within different base composition contexts. There has been continued research into the intimate relationship between the actual DNA sequences, the epigenetic layer of regulatory control, replication timing, and the 2D and 3D genome organisation [4, 46, 54, 2]; such overarching studies may truly benefit from having a general segmentation algorithm such as oHMMed that can be applied to all features of interest and facilitate interpretation of their potential interactions.

Obviously, application of oHMMed is not restricted to genomic data: Any time series data that exhibit the appropriate autocorrelation pattern and conform to the required overall emission distribution can efficiently be segmented into statistically distinct regions using oHMMed; other fields of application may include ecology or indeed also econometrics. The oHHMed algorithms themselves can be extended to include other convex emission densities.

## 5. Supplementary Information

### 5.1. Additional File 1

Mathematical details on the MCMC sampler underlying the oHMMed algorithms.

### 5.2. Additional File 2

Full results of application of oHMMed to the GC proportion and gene content of mouse and fruit fly genomes.

### 5.3. Additional File 3

Full results of the comparison between oHMMed and the equivalent unordered algorithms on sequences simulated with oHMMed and the human genomic data.

### 5.4. Additional File 4

Comparison between oHMMed and the equivalent unordered algorithm on sequences simulated with the latter.

### 5.5. Additional File 5

Extensive documentation of the results of application of oHMMed to the epigenetic markers ATAC, H3K27ac, and H3K27me3 in both 100 *kb* and 1*kb* windows along the human chromosome 1.

## Supporting information

AdditionalFile1

AdditionalFile2

AdditionalFile3

AdditionalFile4

AdditionalFile5

## 6. Declarations

### 6.1. Ethics Approval and Consent to Participate

Not applicable

### 6.2. Consent for Publication

Not applicable

### 6.3. Availability of Data and Materials

The algorithms presented here have been implemented in the R package oHMMed, which is available on CRAN[38], and further information on the package is available on GitHub (https://github.com/LynetteCaitlin/oHMMed)[39]. The raw data used in this article is available at the cited sites within the main text. The processed data, augmented with the corresponding oHMMed-inferred hidden states, is available on GitHub[39] in the files “GenomeAnnotations.zip” and “EpiGenomeAnnotation.txt” in the folder “Data”, and sketches for how to obtain all the analyses and some of the figures presented here can be found in the file “oHMMedOutputAnalyses.R” in the folder “simulation scripts”.

### 6.6. Author Contributions

Concept: CV, CK, JB, LCM, MK;

Planning, Coordination: LCM, CV;

Method - Theory, Usage, Interpretation: LCM, CV;

Method - Implementation, Debugging: CV, LCM,

contributions by MK;

Method - Optimisation, Package Development: MM;

Method - Comparisons: LCM, MK, CV;

Data Handling and Analyses: LCM, BY, JB, HK, CV, MK;

Writing: LCM, CV,

contributions by MK;

Editing and Manuscript Approval: All authors.

## 6.7. Acknowledgements

Not applicable

