## AdditionalFile1 for "Inference of Genomic Landscapes using Ordered Hidden Markov Models with Emission Densities (oHMMed)"

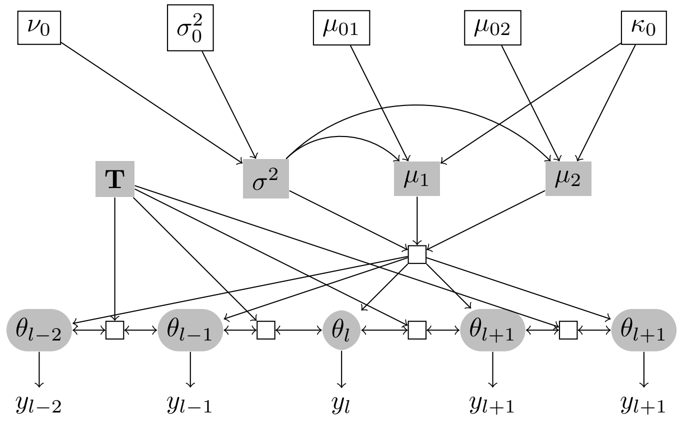


Figure 1: Graphical representation of oHMMed with normal emission densities: Starting at the top of the graph, the hyperparameters from which the emitted densities are drawn must be set: The variance $\sigma^2$ is assumed to be drawn from a prior inverse chi-square distribution with prior degrees of freedom $\nu_0$ and prior variance $\sigma_0^2$; the means are assumed to be drawn from a normal distribution with variance $\sigma^2$ (since we set the coupling constant $\kappa_0 = 1$) and prior means $\boldsymbol{\mu}_0$. The priors for the transition matrix $\mathbf{T}$ are omitted here. The sequence of hidden states $\theta_l$ near the bottom of the graph each 'select' a $\mu_i$, and conditional on the chosen $\mu_i$ and $\sigma^2$, the emitted/observed sequence of data points are drawn from the respective normal distribution.

### 1. Section A1: oHMMed with Normal Emissions

Here we provide a full description of oHMMed with normal emission densities (Fig 1).

Assume that each of the $K$ hidden states emits a normal distribution; traversing the genome, realisations $y_l$ of these normal distributions are observed at each position without knowledge about which is the underlying state. Denote the vector of means per state as $\boldsymbol{\mu} = (\mu_1, \ldots, \mu_K)$, and note the entries are ordered by increasing size. Further, denote the shared standard deviation as $\sigma$. Let us first review how the observed sequence is annotated, *i.e.,* how at each position the posterior probability of the hidden states are calculated via the classic forward-backward algorithm [*e.g.,* 1]. We then provide a Markov chain Monte Carlo algorithm using sampling from the conditional distributions, *i.e.,* a Gibbs sampler [2, chapter 11.3].

#### *1.1. Forward-backward HMM Algorithm*

Define $L \times K$ matrices of auxiliary forward variables $\mathbf{F}$ and backward variables $\mathbf{B}$, with the row-index $l$ defining the position along the genome and column-index $i$ the assigned state. Introduce a vector of $L$ weights $\boldsymbol{s}$. Then, for each position $l$, define $\mathbf{D}_l$ as the diagonal matrix of emission likelihoods:

$$\mathbf{D}_l = \text{diag}\left(\Pr(y_l \,|\, \mu_1, \sigma), \ldots, \Pr(y_l \,|\, \mu_K, \sigma)\right). \tag{1}$$

Denoting a column vector of ones with $\mathbf{1'}$, the forward algorithm can be written as:

- *Initialization ($l = 1$):* Set $s_1 = \boldsymbol{\pi}\mathbf{D}_1\mathbf{1}^{'}$ and $\mathbf{F}_{1\cdot} = \boldsymbol{\pi}\mathbf{D_1}/s_1$
- *Recursion ($l = 2, \ldots, L-1$):* Calculate $s_{l+1} = \mathbf{F}_{l\cdot}\mathbf{T}\mathbf{D}_{l+1}\mathbf{1}^{'}$ and $\mathbf{F}_{(l+1)\cdot} = \mathbf{F}_{l\cdot}\mathbf{T}\mathbf{D}_{l+1}/s_{l+1}$.
- *Stopping rule:* Stop when $\mathbf{F}_{L\cdot}$ is reached.

The corresponding backward algorithm is:

- *Initialization ($l = L$):* Set $\mathbf{B}^{'}_{L\cdot} = \mathbf{1'}/s_L$.
- *Recursion ($l = L-1, \ldots, 1$):* Calculate $\mathbf{B}^{'}_{l\cdot} = \mathbf{T}\mathbf{D}_{l+1}\mathbf{B}^{'}_{(l+1)\cdot}/s_l$.
- *Stopping rule:* Stop when $\mathbf{B}_{1\cdot}$ is reached.

Note that $s_1 = \Pr(y_1 \,|\, \mathbf{T}, \boldsymbol{\mu}, \sigma)$ and $s_l = \Pr(y_l \,|\, y_{l-1}, \mathbf{T}, \boldsymbol{\mu}, \sigma)$ for $2 \leq l \leq L$. It follows that the likelihood is $\Pr(\mathbf{y} \,|\, \mathbf{T}, \boldsymbol{\mu}, \sigma) = \prod_{l=1}^{L} s_l$. But calculating the likelihood in this way may result in underflow with double precision floats and therefore we instead calculate the log likelihood: $\log(\Pr(\mathbf{y}|\mathbf{T}, \boldsymbol{\mu}, \sigma)) = \sum_{l=1}^{L} \log(s_l)$. Further, note that $\mathbf{F}_{l\cdot} = \Pr(\theta_l; y_1, \ldots, y_l \,|\, \mathbf{T}, \boldsymbol{\mu}, \sigma)/(\prod_{k=1}^{l} s_k)$ and $\mathbf{B}_{l\cdot} = \Pr(y_{i+1}, \ldots, y_L \,|\, \theta_i, \mathbf{T}, \boldsymbol{\mu}, \sigma)/\prod_{k=l}^{L} s_k$.

The probability of $\theta_l$ being in state $i$ is:

$$\Pr(\theta_l = i \,|\, y, \mathbf{T}, \boldsymbol{\mu}, \sigma) = \mathbf{F}_{li}\mathbf{B}_{li}s_l\,. \tag{2}$$

The joint probability of two subsequent states $\theta_l = i$ and $\theta_{l+1} = j$ is:

$$\Pr(\theta_l = i, \theta_{l+1} = j \,|\, \boldsymbol{y}, \mathbf{T}, \boldsymbol{\mu}, \sigma) = F_{li}\mathbf{T}_{ij}\mathbf{D}_{(l+1)jj}\mathbf{B}^{'}_{(l+1)j}\,. \tag{3}$$

*1.2. Markov Chain Monte Carlo algorithm*

Here, we wrap the HMM framework in an MCMC sampler.

*Priors.* The user-specified vector of prior means, $\boldsymbol{\mu}_0$ (indexed by $i \,\epsilon\, (1, \ldots, K)$), and the user-specified prior variance, $\sigma_0^2$, specify a proper inverse chi-squared-normal prior distribution

$$\Pr(\boldsymbol{\mu}, \sigma^2 \,|\, \boldsymbol{\mu_0}, \kappa_0, \nu_0, \sigma_0^2) = \Pr(\sigma^2 \,|\, \nu_0, \sigma_0^2) \prod_{i=1}^{K} \Pr(\mu_i \,|\, \mu_{0i}, \sigma^2/\kappa_0)$$

with:

$$\begin{aligned} \Pr(\sigma^2 \,|\, \nu_0, \sigma_0^2) &= \frac{(\nu_0/2)^{\nu_0/2}}{\Gamma(\nu_0/2)}\, \sigma_0^{\nu_0} (\sigma^2)^{-\frac{\nu_0}{2}-1} e^{-\frac{\nu_0\sigma_0^2}{2\sigma^2}} \\ \Pr(\mu_i \,|\, \mu_{0i}, \sigma^2/\kappa_0) &= \sqrt{\frac{\kappa_0}{2\pi\sigma^2}}\, e^{-\frac{\kappa_0(\mu_i - \mu_{0i})^2}{2\sigma^2}}\;, \end{aligned} \tag{4}$$

where $\nu_0$ represents the prior degrees of freedom and $\kappa_0$ is a coupling constant (which can both generally be chosen freely, but are set to one in this implementation). Note that setting the coupling constant to one simplifies setting "weakly informative" priors.

*Iterations.*

- *Initialise:* An initial transition matrix $\mathbf{T}^{(0)}$, an initial emission vector $\boldsymbol{\mu}^{(0)}$, and an initial standard deviation $\sigma^{(0)}$ are set by the user. Alternatively, $\mathbf{T}^{(0)}$ is set to the user specified prior and the other initial variables are sampled from the prior distributions using priors set by the user as shown above. Conditional on the transition matrix, the stationary distribution, *i.e.,* the row vector $\boldsymbol{\pi}^{(0)}$, is calculated. Set the iteration index $t = 1$, and pick the maximum number of iterations $t = T$.

- *Step 1: Updating $\theta$:* For $t \,\epsilon\, (1, \ldots, T)$, calculate the probability of the state at the first position on the sequence as:

$$\Pr(\theta_1^{(t)} \,|\, \boldsymbol{y}, \mathbf{T}^{(t-1)}, \boldsymbol{\mu}^{(t-1)}, \sigma^{(t-1)}, \boldsymbol{\pi}^{(t-1)}) = \mathbf{F}_{1i}^{(t-1)} \mathbf{B}_{1i}^{(t-1)} s_1^{(t-1)} \,.$$

Sample the state $\theta_1^{(t)}$ from this probability vector, and calculate all further states $\theta_l^{(t)}$ using the above forward-backward HMM algorithm by recursively drawing $\theta_l$ conditional on $\theta_{l-1} = i$ and the other variables using:

$$\Pr(\theta_{l+1}^{(t)} = j \,|\, \theta_l^{(t)} = i, y, \mathbf{T}^{(t-1)}, \boldsymbol{\mu}^{(t-1)}, \sigma^{(t-1)}) = \frac{\mathbf{T}_{ij}^{(t-1)} \mathbf{D}_{(l+1)jj}^{(t-1)} \mathbf{B}_{(l+1)j}^{\prime(t-1)}}{\sum_j \mathbf{T}_{ij}^{(t-1)} \mathbf{D}_{(l+1)jj}^{(t-1)} \mathbf{B}_{(l+1)j}^{\prime(t-1)}} \,. \tag{5}$$

This completes the annotation at time $t$. The marginal log-likelihood

$$lh^{(t-1)} = \sum_{l=1}^{L} \log\left(s_l^{(t-1)}\right)$$

is easily determined at this stage.

- *Step 2: Updating the means $\boldsymbol{\mu}$ and the standard deviation $\sigma$:* Conditional on the current annotation, $\theta_1^{(t)}, \ldots, \theta_L^{(t)}$, calculate the empirical means and standard deviations. Introduce the indicator variable $\mathbf{1}_{\theta_l^{(t)}=i}$, which is one if $\theta_l^{(t)} = i$ and zero otherwise. Then calculate

$$L_i^{(t)} = \sum_l \mathbf{1}_{\theta_l^{(t)}=i} \,,$$

$$\bar{y}_i^{(t)} = \frac{\sum_l \mathbf{1}_{\theta_l^{(t)}=i} \big(y_l^{(t)}\big)}{L_i^{(t)}}$$

and

$$SS^{(t)} = \sum_i \sum_l \mathbf{1}_{\theta_l^{(t)}=i} \big(y_l^{(t)} - \bar{y_i}^{(t)}\big)^2 \,.$$

Sample the standard deviation $\sigma^{(t)}$ from an inverse scaled chi-squared distribution [compare 2, chapter 3.3]

$$\sigma^{2^{(t)}} \sim \text{Inv-}\chi^2(\nu_p, \sigma_p^{2^{(t)}}) \,.$$

with posterior degrees of freedom $\nu_p = 1+L$ and posterior scale parameter

$$\sigma_p^{2(t)} = \left( SS^{(t)} + \sigma_0^2 + \sum_i \left( \left(\mu_{0i} - \bar{y}_i^{(t)}\right)^2 \frac{L_i}{1+L_i} \right) \right) / (1+L)$$

Sample the new vector of means from a normal distribution [compare 2, chapter 3.3] with posterior coupling factors $\kappa_{pi} = 1 + L_i^{(t)}$ and posterior means $\mu_{pi}^{(t)} = \frac{\mu_{0i}+L_i^{(t)}\bar{y}_i^{(t)}}{1+L_i^{(t)}}$ for $i$ in $1 \le i \le K$:

$$\mu_i^{(t)} \sim \mathrm{N}(\mu_{pi}^{(t)}, \sigma^{(t)}/\kappa_{pi}) \,.$$

Then sort the newly sampled means to satisfy the constraint $\mu_i^{(t)} \le \mu_j^{(t)}$ for $i < j$.

- *Step 3: Updating* $\mathbf{T}$*:*Given $\theta_l^{(t)}$, count the pairs $C_{ij}^{(t)} = \sum_l (\theta_l^{(t)} = i, \theta_{l+1}^{(t)} = j)$. Next, sample an auxiliary vector with $2K-2$ entries from a Dirichlet distribution with following parameters: The first $K$ Dirichlet parameters are $\alpha_{ii} + C_{ii}^{(t)}$, and the next $K-1$ are $\alpha_{ij} + \alpha_{ji} + C_{ij}^{(t)} + C_{ji}^{(t)}$, where pseudo-counts $\alpha_{ij}$, corresponding to the prior flux $\mathrm{diag}\,(\boldsymbol{\pi}_0)\mathbf{T}_0$, are added as proper priors. Let the $K \times K$ matrix $\mathbf{P}$ represent the joint probabilities of $\theta_l$ and $\theta_l + 1$. Recall that we assume reversibility, *i.e.,* symmetry of $\mathbf{P}$ conditionally on the states $\theta_l$ and $\theta_{l+1}$. Enter the first $K$ samples of the auxiliary vector on the main diagonal of $\mathbf{P}$; divide the next $K-1$ samples by two and enter them on the diagonals below and above the main diagonal of $\mathbf{P}$; set all other positions of $\mathbf{P}$ to zero. Obtain the new transition matrix $\mathbf{T}^{(t)}$ by normalizing the rows of $\mathbf{P}$, and a new $\boldsymbol{\pi}^{(t)}$ as the stationary distribution of $\mathbf{T}^{(t)}$.
- *IF* $t+1 < T$*:* Set $t$ to $t+1$ and return to *Step 1*
- *IF* $t = T$*:* Stop.

*Estimates.* The most likely hidden state $\widehat{\theta_l}$ for each position along the observed sequence is determined as the highest marginal probability of states $1,..,K$ at that position averaged over the output of *Step 1* from the end of the burn-in period (which is either user specified or left at the algorithm's internally set value) up to $t = T$. The final estimates of $\hat{\mathbf{T}}$, $\hat{\boldsymbol{\mu}} = (\widehat{\mu_1}, \ldots, \widehat{\mu_K})$, and $\hat{\sigma}$ are determined as averages of the sequence of posterior estimates generated by *Step 2*, *Step 3* from the end of the burn-in period up to $t = T$.

*Model fit.* Checks for convergence and assessment of the final model fit are provided in the R package oHMMed and described in the usage recommendations on GitHub [3].

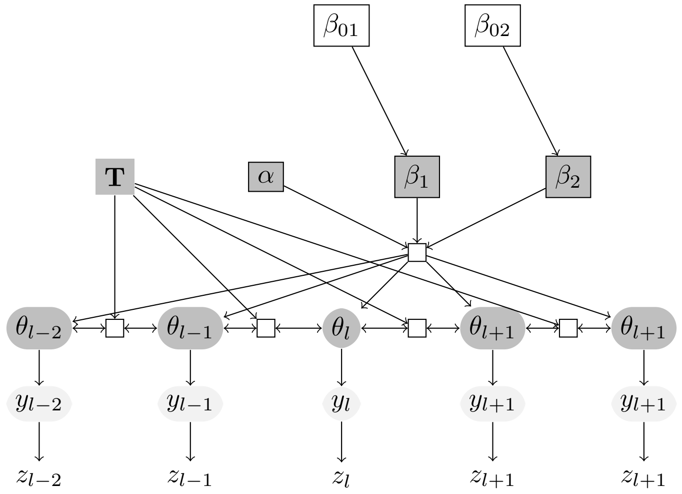


Figure 2: Graphical representation of oHMMed with gamma-poisson emission densities: Starting at the top of the graph, the hyperparameters from which the emitted densities are drawn must be set: The rate parameters $\beta_i$ are assumed to be drawn from exponential distributions with priors $\beta_{0i}$. The improper prior for the shape parameter $\alpha$ cannot be properly shown and the priors for the transition matrix $\mathbf{T}$ are omitted.The sequence of hidden states $\theta_l$ near the bottom of the graph each 'select' a $\beta_i$, and conditional on the chosen $\beta_i$ and $\alpha$, the emitted sequence of data points $y_l$ are drawn from the gamma normal distribution. These are rate parameters, which are used to draw the sequence of poisson distributed observed data points $z_l$.

### 2. Section A2: oHMMed with Gamma-Poisson Emissions

Here we provide a description of oHMMed with gamma-poisson emission densities (Fig 2). We will utilise the backbone of the methodology outlined in the full description of oHMMed with normal emissions (Supplementary Data A1 (1)), and only detail the required changes.

#### *2.1. From Emissions of Auxiliary Variables to Observations*

Assume that the observed sequence consists of count data, and that the overall distribution of these counts resembles a mixture of poisson distributions. Therefore, at each position $l$ along the chromosome, the probability of each realised count $z_l$ is:

$$\Pr(z_l \,|\, y_l) = \frac{y_l^{z_l} e^{-y_l}}{z_l!} \,. \tag{6}$$

The emitted auxiliary variables $y_l$ are assumed to be drawn from a mixture of gamma distributions with common shape parameter parameter $\alpha$ and state-specific rate parameters $\beta_i$:

$$\Pr(y_l \,|\, \alpha, \beta_i, \theta_l = i) = \frac{\beta_i^{\alpha}}{\Gamma(\alpha)}\, y_l^{\alpha-1} e^{-\beta_i y_l} \,, \tag{7}$$

where $0 < \alpha, \beta_i < \infty$. Marginalizing, by integrating over the auxiliary variables, provides the likelihoods at specific positions needed for $\mathbf{D}_l$ (compare Eq. 1) in the forward-backward algorithm:

$$\begin{aligned}
\Pr(z_l \mid \alpha, \beta_i) &= \int_0^\infty \Pr(z_l \mid y_l) \Pr(y_l \mid \alpha, \beta_i, \theta_l = i)\, dy_l \\
&= \int_0^\infty \frac{y_l^{z_l} e^{-y_l}}{z_l!} \frac{\beta_i^\alpha}{\Gamma(\alpha)}\, y_l^{\alpha-1} e^{-\beta_i y_l}\, dy_l \\
&= \frac{1}{z_l!} \frac{\beta_i^\alpha}{\Gamma(\alpha)} \int_0^\infty y_l^{\alpha+z_l-1} e^{-(\beta_i+1)y_l}\, dy_l \\
&= \frac{\Gamma(\alpha+z_l)}{z_l!\,\Gamma(\alpha)} \big(\tfrac{\beta_i}{\beta_i+1}\big)^\alpha \big(\tfrac{1}{\beta_i+1}\big)^{z_l} .
\end{aligned} \tag{8}$$

We note that the last line is a parametrization of the negative binomial distribution, while the integrand in the third line will be important for updating during the MCMC sampler immediately below.

*2.2. MCMC Sampler*

Here, we provide the priors and show how to update the parameters of the emitted gamma distributions as part of adapting *Step 2* from the MCMC sampler in Supplementary Data A1 (1.2) to the scenario of gamma-poisson emissions.

*Priors.* For the parameters $\alpha$ and the vector $\boldsymbol{\beta} = (\beta_1, \ldots, \beta_K)$ of the gamma distributions, semi-conjugate or conditionally conjugate priors are assumed, *i.e.,* the joint prior distribution does not factorise [2]. In particular, for each $\beta_i$ a prior gamma distribution with shape parameter $\delta_0$ and rate parameter $\delta_0/\beta_{0i}$ is assumed:

$$\Pr(\beta_i \mid \delta_0, \beta_{0i}) \sim \text{Gamma}(\delta_0, \delta_0/\beta_{0i}) = \frac{(\delta_0/\beta_{0i})^{\delta_0}}{\Gamma(\delta_0)} (\beta_i)^{\delta_0 - 1} e^{\frac{\delta_0 \beta_i}{\beta_{0i}}} . \tag{9}$$

We note that with this parametrization, the expectation of $\beta_i$ is $\beta_{0i}$, which we call the "prior beta" when setting "weakly informative" priors usage recommendations on GitHub [3]. In our calculations, we assume $\delta_0 = 1$, resulting in an exponential prior distribution. The conditional conjugate prior of $\alpha$ is not standard:

$$\Pr(\alpha \mid \upsilon_0, \lambda_0) = \frac{1}{\int_0^\infty \big(\frac{\upsilon_0^{\alpha-1}}{\Gamma(\alpha)}\big)^{\lambda_0} d\alpha} \left(\frac{\upsilon_0^{\alpha-1}}{\Gamma(\alpha)}\right)^{\lambda_0} , \tag{10}$$

where the first term to the right is the constant of proportionality, which we will denote with $c$ further down. While integration of the function (10) to obtain $c$ may be difficult, it is not needed for updating the MCMC sampler, as shown below. For $\lambda_0 = 1$, the distribution (10) with the single parameter $\upsilon_0$ can be thought of as a continuous generalization of the discrete Poisson distribution, since the Gamma function is the continuous generalization of the factorial. For increasing $\upsilon_0$ and setting $i = \alpha + 1$, we have for $\lambda_0 = 1$:

$$
\begin{aligned}
c^{-1} &= \int_0^\infty \frac{\upsilon_0^{\alpha-1}}{\Gamma(\alpha)}\, d\alpha \\
&\approx \sum_{\alpha=0}^{\infty} \frac{\upsilon_0^{\alpha-1}}{\Gamma(\alpha)} \\
&= \frac{\upsilon_0^{-1}}{\Gamma(0)} + \sum_{i=0}^{\infty} \frac{\upsilon_0^{i}}{i!} \\
&= \upsilon_0^{-1} + e^{\upsilon_0} \sum_{i=0}^{\infty} \frac{\upsilon_0^{i} e^{-\upsilon_0}}{i!} \\
&= \upsilon_0^{-1} + e^{\upsilon_0}\,,
\end{aligned}
\tag{11}
$$

where the second line follows from the first as the integral converges to the corresponding sum with increasing grid points and the last line follows from the previous by noting that the sum over the Poisson distribution function is one. For large $\upsilon_0$, it thus follows that $c \approx e^{-\upsilon_0}$ and the similarity to the Poisson distribution is obvious. Furthermore

$$
\begin{aligned}
\mathrm{E}(\alpha \,|\, \upsilon_0) + 1 &= c \int_0^\infty (\alpha+1) \left( \frac{\upsilon_0^{\alpha-1}}{\Gamma(\alpha)} \right) d\alpha \\
&= c \int_1^\infty i \left( \frac{\upsilon_0^{i}}{\Gamma(i+1)} \right) di \\
&\approx c \sum_{i=1}^{\infty} i \frac{\upsilon_0^{i}}{i!} \\
&= c e^{\upsilon_0} \sum_{i=1}^{\infty} i \, \frac{\upsilon_0^{i} e^{-\upsilon_0}}{i!} \\
&= c e^{\upsilon_0} \left( -e^{-\upsilon_0} + \sum_{i=0}^{\infty} i \, \frac{\upsilon_0^{i} e^{-\upsilon_0}}{i!} \right) \\
&= c e^{\upsilon_0} \big( -e^{-\upsilon_0} + \upsilon_0 \big)\,,
\end{aligned}
\tag{12}
$$

where the third line follows from the second as the integral converges to the corresponding sum with increasing grid points and the last line follows from the previous as the expectation of the Poisson distribution is its parameter. Hence for large $\upsilon_0$, $c \approx e^{-\upsilon_0}$ and $\mathrm{E}(\alpha \,|\, \upsilon_0) \approx \upsilon_0 - 1$.

For very low $\upsilon_0$, the mean approaches 0 (see Fig. 3). Note that for $\upsilon_0 \geq 1$, the mean is close to $\upsilon_0 + 1$ and the mode to $\upsilon_0 + 1/2$. The mean and variance of the Poisson distribution are both equal to the rate parameter; the product of $L$ such distribution functions results in a distribution with the mean converging to the mode of the individual distribution and $1/L$ times its variance. This can be used to set weakly informative priors (see the usage recommendations on

GitHub[3] ), and for choosing a proposal or "jump" distribution for the MCMC sampler (see below).

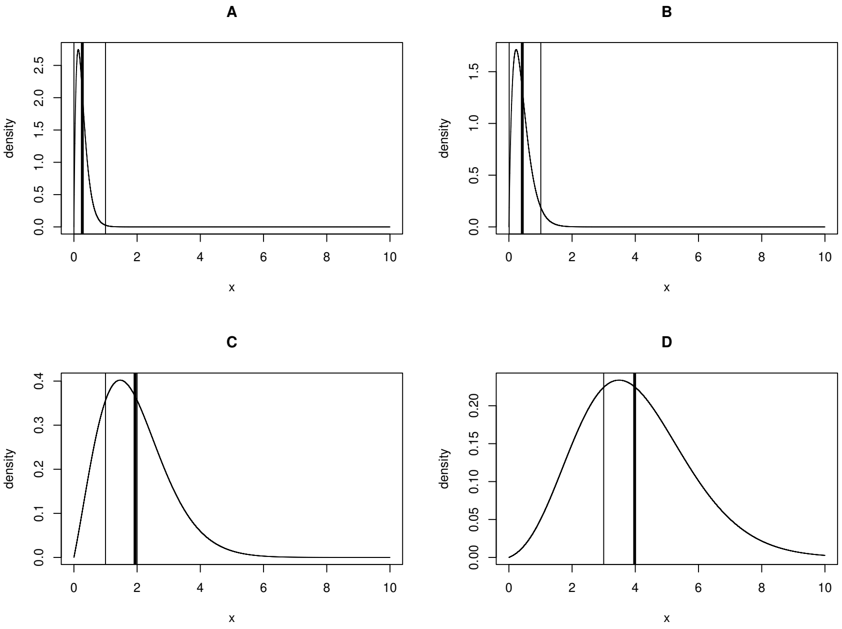


Figure 3: The prior distribution of $\alpha$, assuming $\lambda_0 = 1$ for A) $\upsilon_0 = 0.0005$; B) $\upsilon_0 = 0.01$; C) $\upsilon_0 = 1$; D) $\upsilon_0 = 3$. The thick line corresponds to the mean of the distribution; the left thin line to $\upsilon_0$, the right thin line to $\upsilon_0 + 1$. Note that mean and $\upsilon_0 + 1$ nearly coincide for $\upsilon_0 \geq 1$.

*Updating $y_l$.* The starting point is the posterior conditional distribution of an emitted auxiliary variable $y_l$ at position $l$ along the chromosome; note that we assume the current $\alpha^{(t)}$ and $\beta_i^{(t)}$ and a current annotation $\theta_l^{(t)} = i$ to be available. We sample the auxiliary variables $y_l^{(t)}$ from their conditional posterior (compare the third line in Eq 8):

$$\Pr(y_l \,|\, z_l, \alpha^{(t)}, \beta_i^{(t)}) = \frac{(\beta_i^{(t)} + 1)^{\alpha^{(t)} + z_l}}{\Gamma(\alpha^{(t)})} y_l^{\alpha^{(t)} + z_l - 1} e^{-(\beta_i^{(t)} + 1) y_l} \,. \tag{13}$$

*Updating $\beta_i$.* Conditional on the full sequence of auxiliary variables $\boldsymbol{y}$ and states $\boldsymbol{\theta}$, $\alpha$ and the $\beta_i$ are independent of the observed sequence data $\mathbf{z}$. We therefore have the marginal likelihood:

$$
\begin{aligned}
\Pr(\boldsymbol{y} \mid \alpha, \beta_i, \theta_l) &= \prod_{l=1}^{N} \mathbf{1}_{\theta_l = i} \Big( \Pr(y_l \mid \alpha, \beta_i, \theta_l) \Big) \\
&= \prod_{l=1}^{N} \mathbf{1}_{\theta_l = i} \left( \frac{\beta_i^{\alpha}}{\Gamma(\alpha)} \, y_l^{\alpha-1} e^{-\beta_i y_l} \right) \\
&= \prod_{i=1}^{K} \mathbf{1}_{\theta_l = i} \left( \prod_{l=1}^{L_i} \frac{\beta_i^{\alpha}}{\Gamma(\alpha)} \, y_l^{\alpha-1} e^{-\beta_i y_l} \right)
\end{aligned} \tag{14}
$$

We note that this function conveniently allows for decomposition with respect to the $\beta_i$, as the terms in the parentheses have the functional form of a gamma distribution. Hence, in a Gibbs sampling step [2, chapter 11.3], each $\beta_i$ can be sampled from a gamma distribution assuming the conjugate prior (9). Conditional on the hidden states, set

$$
\bar{y}_i = \frac{\sum_l \mathbf{1}_{\theta_l^{(t)} = i} y_l}{L_i} \,,
$$

where $\mathbf{1}_{\theta_l = i}$ is an indicator variable that is one if $\theta_l = i$ and zero otherwise and $L_i = \sum_l \mathbf{1}_{\theta_l = i}$. This results in a posterior beta with posterior $\delta_p = \alpha L_i^{(t)} + \delta_0$ and posterior rate parameter

$$
\frac{\delta_p}{\beta_{pi}} = L_i^{(t)} \bar{y}_i^{(t)} + \delta_0 / \beta_{0i} \,.
$$

Then sort the newly sampled rate parameters to satisfy the constraint $\beta_i^{(t)} \geq \beta_j^{(t)}$ for $i < j$.

*Updating $\alpha$.* We use a Metropolis-Hastings sampler [2, chapter 11.2] for updating $\alpha$. With the proper conjugate prior (10), the conditional posterior of $\alpha$ is proportional to:

$$
\begin{aligned}
\Pr(\alpha \mid \boldsymbol{\theta}, \boldsymbol{y}, \beta_i^{(t+1)}) &\propto \left( \frac{\upsilon_0^{\alpha-1}}{\Gamma(\alpha)} \right)^{\lambda_0} \frac{\left( \prod_{l=1}^{L} \mathbf{1}_{y_l = i} \big( \beta_i^{(t+1)} y_l \big) \right)^{\alpha}}{\Gamma(\alpha)^L} \\
&\propto \left( \frac{\upsilon_p^{\alpha-1}}{\Gamma(\alpha)} \right)^{\lambda_p} ,
\end{aligned} \tag{15}
$$

with

$$
\lambda_p = \lambda_0 + L \,.
$$

and

$$
\upsilon_p = \left( \upsilon_0^{\lambda_0} \prod_{l=1}^{L} \mathbf{1}_{y_l = i} \big( \beta_i^{(t+1)} y_l \big) \right)^{\frac{1}{\lambda_0 + L}} .
$$

Note that $\upsilon_p$ is a weighted geometric mean of the prior and the data. As a Metropolis-Hastings proposal or “jump” distribution to sample a proposal value

$\alpha^*$ [2, chapter 11.2] approximating the conditional posterior (15), we choose the gamma distribution:

$$J(\alpha^* \mid \alpha^{(t)}, \boldsymbol{\theta}, \boldsymbol{y}, \beta_i^{(t+1)}) \sim \text{Gamma}(\lambda_p \alpha^{(t)}, \lambda_p)\,.$$

Given a new $\alpha^*$ sampled from this jump distribution, we calculate the decision-rule ratio:

$$r = \frac{\Pr(\alpha^* \mid \boldsymbol{\theta}, \boldsymbol{y}, \beta_i^{(t)}) / J(\alpha^* \mid \alpha^{(t)}, \boldsymbol{\theta}, \boldsymbol{y}, \beta_i^{(t)})}{\Pr(\alpha^{(t)} \mid \boldsymbol{\theta}, \boldsymbol{y}, \beta_i^{(t)}) / J(\alpha^{(t)} \mid \alpha^*, \boldsymbol{\theta}, \boldsymbol{y}, \beta_i^{(t)})}\,. \tag{16}$$

The newly sampled $\alpha^*$ is accepted (*i.e.,* setting $\alpha^{(t)} = \alpha^*$) with probability $min(r, 1)$, and rejected (*i.e.,* leaving $\alpha^{(t)} = \alpha^{(t-1)}$) otherwise [2, chapter 11].
