## AdditionalFile2 for "Inference of Genomic Landscapes using Ordered Hidden Markov Models with Emission Densities (oHMMed)"

### 1. Detailed Results on Segmentation of Mouse and Fruit Fly Genomes

#### *1.1. Segmentation of the Mouse Genome*

*GC Proportion.* As in the human case, $K = 5$ hidden states are considered a good fit for the variation in GC proportion along the mouse genome (Fig. 1). This results in states with mean emissions that are almost equally spaced and clearly separated statistically, even by our conservative estimate of $\pm$ the standard deviation (see Fig. 1B). Specifically, the estimates for the respective means are 0.344, 0.383, 0.420, 0.460, 0.507 and the shared standard deviation is estimated at 0.017. The GC proportion is thus generally higher and fluctuates within a narrower range than in humans. The proportion of genomic windows assigned to each of these states are 0.158, 0.223, 0.304, 0.213, 0.102 respectively. We infer a transition matrix between hidden states of

$$\begin{pmatrix} 0.933 & 0.067 & 0 & 0 & 0 \\ 0.047 & 0.846 & 0.106 & 0 & 0 \\ 0 & 0.079 & 0.813 & 0.108 & 0 \\ 0 & 0 & 0.150 & 0.769 & 0.080 \\ 0 & 0 & 0 & 0.167 & 0.833 \end{pmatrix},$$

which translates to an average of 16, 7, 6, 5, 6 consecutive windows being assigned to to the respective states.

*Gene Content.* Next, we focus on the density of protein coding genes along the mouse genome. Our algorithms suggest $K = 4$ hidden states with well separated means (Figs. 2 and 3). These inferred means are 0.173,1.160, 2.588, 5.527, and the corresponding states occur at proportions 0.247, 0.541, 0.172, and 0.039.

The transition rate matrix is estimated as

$$\begin{pmatrix} 0.908 & 0.092 & 0 & 0 \\ 0.044 & 0.932 & 0.024 & 0 \\ 0 & 0.067 & 0.910 & 0.023 \\ 0 & 0 & 0.099 & 0.901 \end{pmatrix},$$

leading to an average of 13,19, 14, 12 sequential windows for the three respective states.

*Correlation.* A clear positive correlation between the segmentation of the mouse genome by GC proportion and gene content can be seen by cross-tabulation (visualisation in Fig. 5B of the main text). The position-specific posterior means of the GC content and gene density once again show a highly significant (p$<$ $2.2e^{-16}$) positive Spearman correlation coefficient, this time of 0.57 (see Fig. 1C). A genomic region of particularly steady high positive correlation is apparent on chromosome 3; some shorter interspersed regions exhibit a similar pattern.

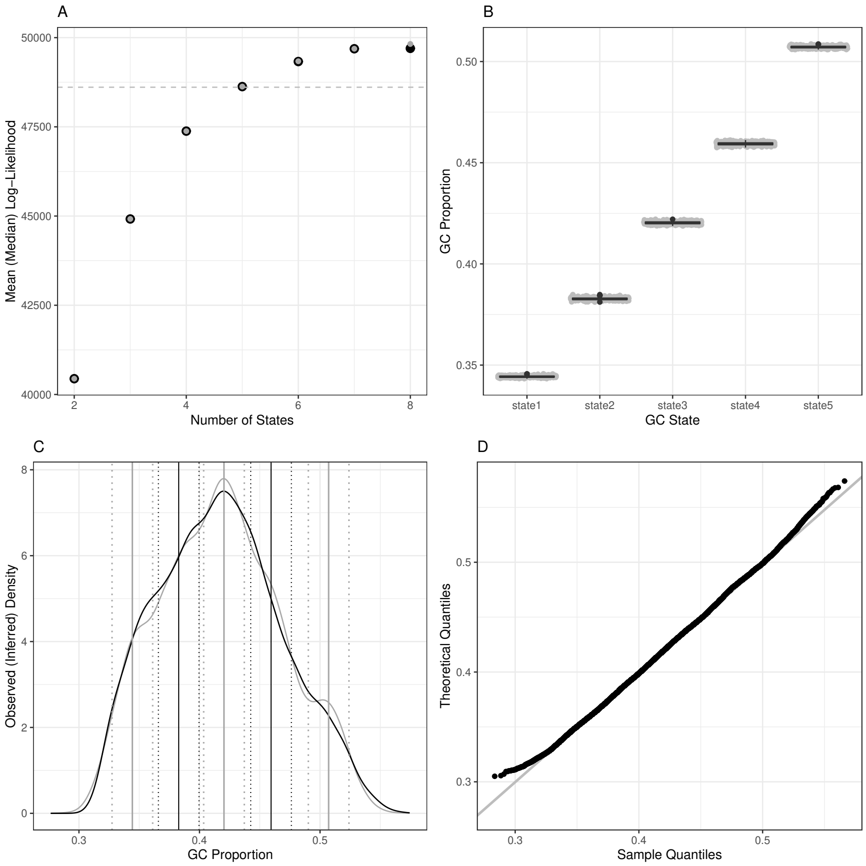


Figure 1: Diagnostics for annotation of the *Mus musculus* genome by average GC proportion using oHMMed with normal emission densities. Panel A shows the mean (black) and median (grey) log-likelihood of fully converged runs of the algorithm with different numbers of hidden states, with the dashed horizontal line marking the selected number of hidden states. In panel B, boxplots of the posterior (i.e. inferred) mean GC proportion of the run with the chosen number of states - which is five - are presented. Panel C shows the observed overall density (black) of the GC proportion superimposed on the posterior (inferred) density, with the inferred means per chosen number of states plus the 68% confidence intervals drawn in vertical lines. The final panel D shows the QQ-plot of the observed density vs. the posterior density (here termed the theoretical distribution).

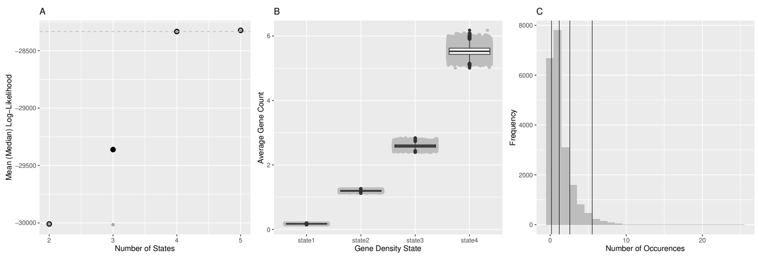


Figure 2: This figure represents the first part of the summarised diagnostics for oHHMed with gamma-poisson emission densities as employed on counts of the average number of protein coding genes along the *Mus musculus* genome. Panel A shows the mean (black) and median (grey) log-likelihood of fully converged runs of the algorithm with different numbers of hidden states, with the dashed horizontal line marking the chosen number - which is four in this case. In panel B, boxplots of the posterior (i.e. inferred) mean gene densities of the inference run with four hidden states are presented. Panel C shows the observed distribution of gene counts with the inferred means superimposed as vertical lines; again, these are significantly different on the 95% confidence level as per one-sided poisson rate test.

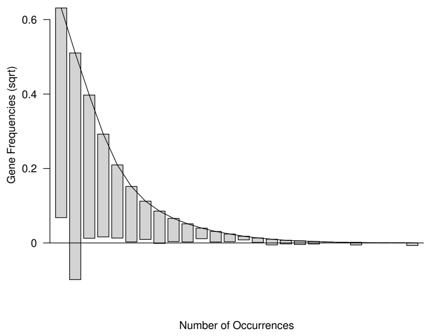


Figure 3: As the final part of the summarised diagnostics for oHMMed with gamma-poisson emission densities and four hidden states as applied to the protein coding genes in *Mus musculus*, we present the above rootogram: The bars represent the observed frequency of counts (square root transformed), and they have been shifted so that the top of each bar aligns with the (smoothed) distribution inferred by oHMMed. Deviations can therefore be assessed by checking the distance of the lower end of each bar to the x-axis.

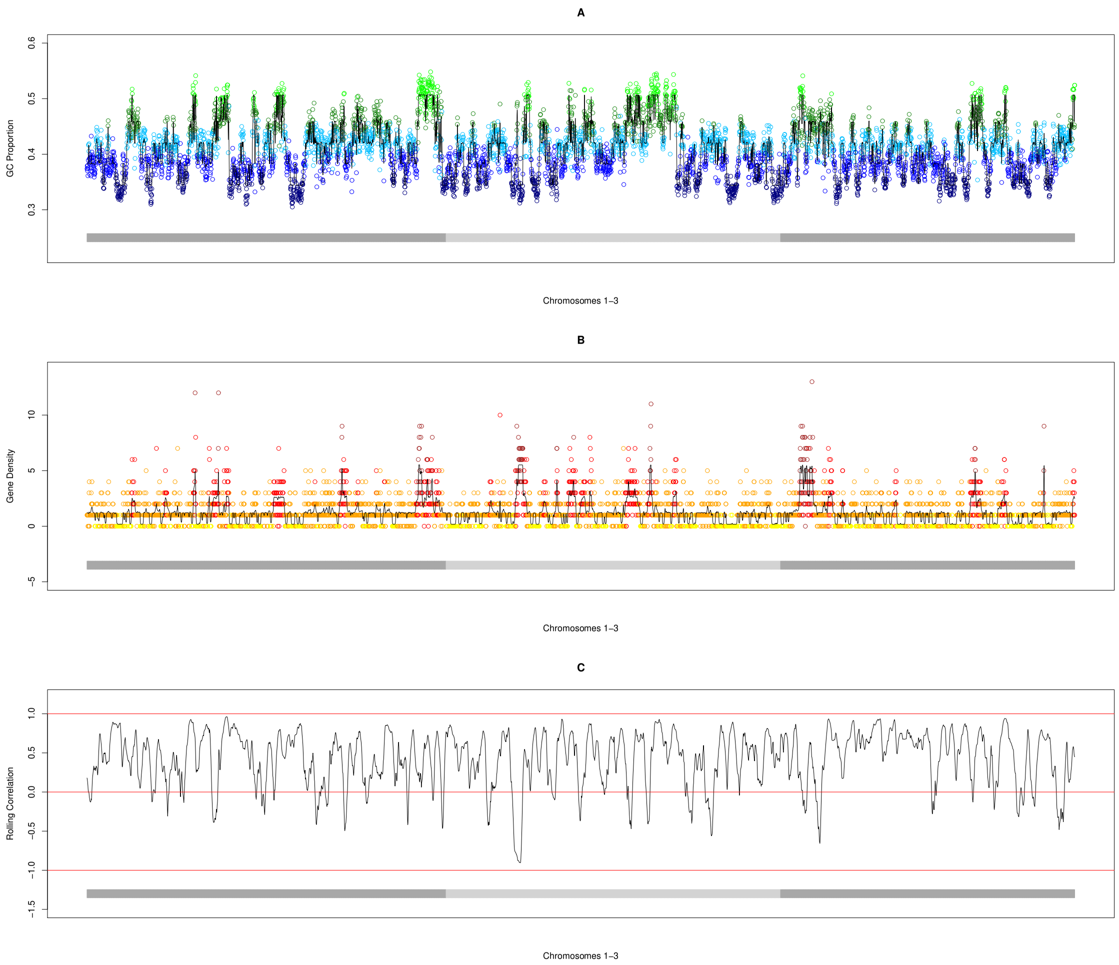


Figure 4: In each of the above panels A-C, the *Mus musculus* chromosomes $1-3$ (demarked by alternating dark and light grey horizontal bars) are plotted for different oHMMed analyses: In A, the average GC proportion in every 100kb window is coloured by the oHMMed-inferred GC state. The three lower states are in blue and the two higher ones in green, with the shades lightening with higher GC proportion. In B, the number of protein coding genes for every 100kb window are shown in colours corresponding to the oHMMed-inferred gene density states: yellow, orange, red, and brown mark increasing gene density states. Note that in both A and B, the black lines trace the posterior (inferred) means returned by oHMMed with normal and gamma-poisson emissions respectively. These position-specific posterior means are the sum of estimated means times the respective probabilities of each state, thus combining both estimated mean values and the algorithm's certainty of the assigned state. In C, Spearman's correlation for the two posterior means in shown in rolling windows of 50 collated 100kb windows.

*1.2. Segmentation of the Fruit Fly Genome*

*GC Proportion.* The GC proportion along the genome of *Drosophila melanogaster* appears fairly uniform in comparison to the other species examined in this paper (Fig. 8A). Nonetheless, we are able to model $K = 4$ hidden states (Fig. 5A) with clearly separated mean emissions of 0.387, 0.412, 0.443, and 0.475 and a standard deviation of 0.022; even so the majority of the genome is actually assigned to states of intermediate GC content with the exact splits according to each state being 0.117, 0.423,0.400, 0.060. Note that this is comparable to the results in [1, Table 1], which further shows that the GC proportion of *D. melanogaster* is relatively low and variable compared to other members of the genus. Our inferred transition rate matrix between hidden states is

$$\begin{pmatrix} 0.929 & 0.071 & 0 & 0 \\ 0.022 & 0.875 & 0.103 & 0 \\ 0 & 0.115 & 0.840 & 0.046 \\ 0 & 0 & 0.255 & 0.745 \end{pmatrix},$$

which results in realised consecutive windows of equal states with average lengths of 18,10, 8, 5 respectively. Note that much of the longer stretches of low GC regions are focused around the telomeric regions (Fig. 8A).

*Gene Content.* The number of protein coding genes per window across the *D. melanogaster* genome is even more homogeneous (Fig. 8B). We infer only $K = 2$ distinct hidden states, present in proportions of 0.553 and 0.447, with means of 0.421 and 2.138. The states are assigned comparable stretches of subsequent windows on average (12 and 10), with transitions:

$$\begin{pmatrix} 0.900 & 0.100 \\ 0.119 & 0.881 \end{pmatrix},$$

For the visual diagnostics that correspond to these results, see Figs. (6 and 7).

*Correlation.* Despite not being able to segment the *D. melanogaster* into as many states according to either the GC proportion or content of protein coding genes as the other species, cross-tabulating these reveals a positive correlation (Fig. 5C in the main text), and also compare [1, Fig. 7]. This is corroborated by the highly significant (p$< 2.2e^{-16}$) positive Spearman correlation coefficient of 0.45 between the position-specific posterior means of the GC content and gene density (see also Fig. 5C).

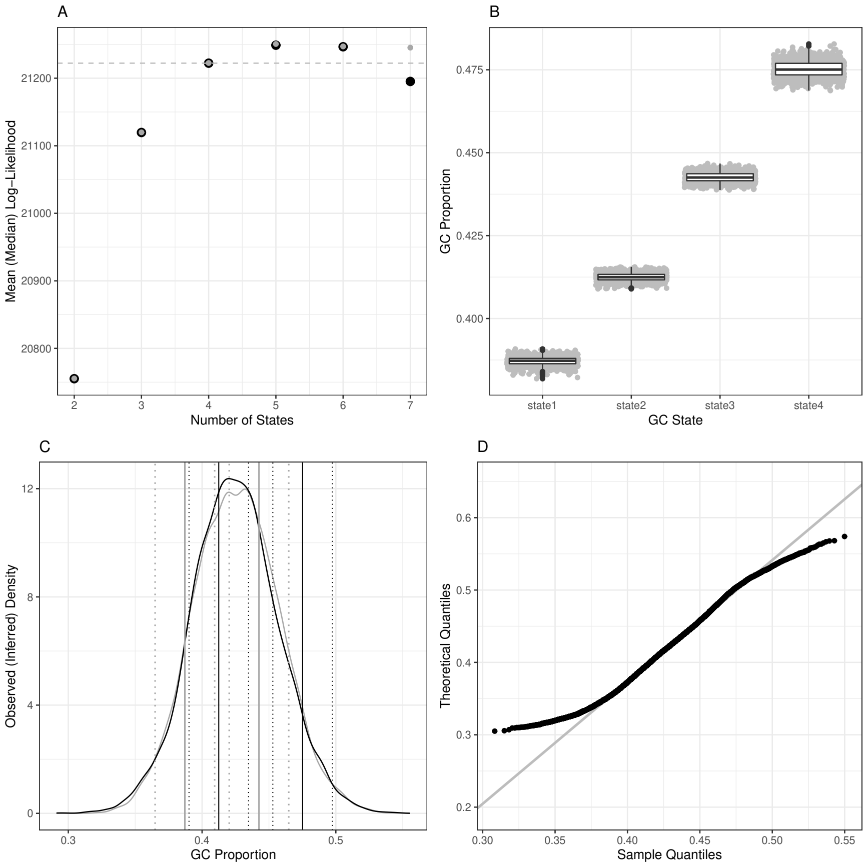


Figure 5: Diagnostics for annotation of the *Drosophila melanogaster* genome by average GC proportion using oHMMed with normal emission densities. Panel A shows the mean (black) and median (grey) log-likelihood of fully converged runs of the algorithm with different numbers of hidden states, with the dashed horizontal line marking the selected number of hidden states. The difference between the mean and median for three hidden states is the effect of autocorrelation in the traces of the estimated parameters. In panel B, boxplots of the posterior (i.e. inferred) mean GC proportion of the run with the chosen number of states - which is four - are presented. Panel C shows the observed overall density (black) of the GC proportion superimposed on the posterior (inferred) density, with the inferred means per chosen number of states plus the 68% confidence intervals drawn in vertical lines. The final panel D shows the QQ-plot of the observed density vs. the posterior density (here termed the theoretical distribution).

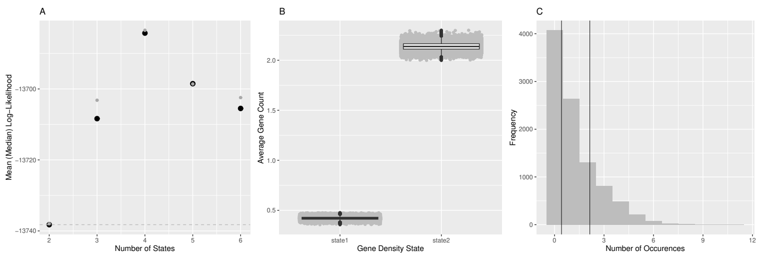


Figure 6: Here we show the first part of the summarised diagnostics for oHHMed with gamma-poisson emission densities as employed on counts of the average number of protein coding genes along the *Drosophila melanogaster* genome. Panel A shows the mean (black) and median (grey) log-likelihood of fully converged runs of the algorithm with different numbers of hidden states, with the dashed horizontal line marking the chosen number - which is only two in this case. Note that the mean and medians often do not coincide here; this is indicative of more erratic autocorrelation in the traces of the estimators, but is not an issue for two hidden states. In panel B, boxplots of the posterior (i.e. inferred) mean gene densities of the inference run with two hidden states are presented. Panel C shows the observed distribution of gene counts with the inferred means superimposed as vertical lines; these are significantly different on the 95% confidence level as per one-sided poisson rate test.

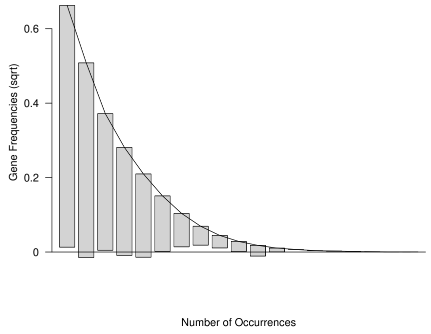


Figure 7: As the final part of the summarised diagnostics for oHMMed with gamma-poisson emission densities and two hidden states as applied to the protein coding genes in *Drosophila melanogaster*, we present the above rootogram: The bars represent the observed frequency of counts (square root transformed), and they have been shifted so that the top of each bar aligns with the (smoothed) distribution inferred by oHMMed. Deviations can therefore be assessed by checking the distance of the lower end of each bar to the x-axis.

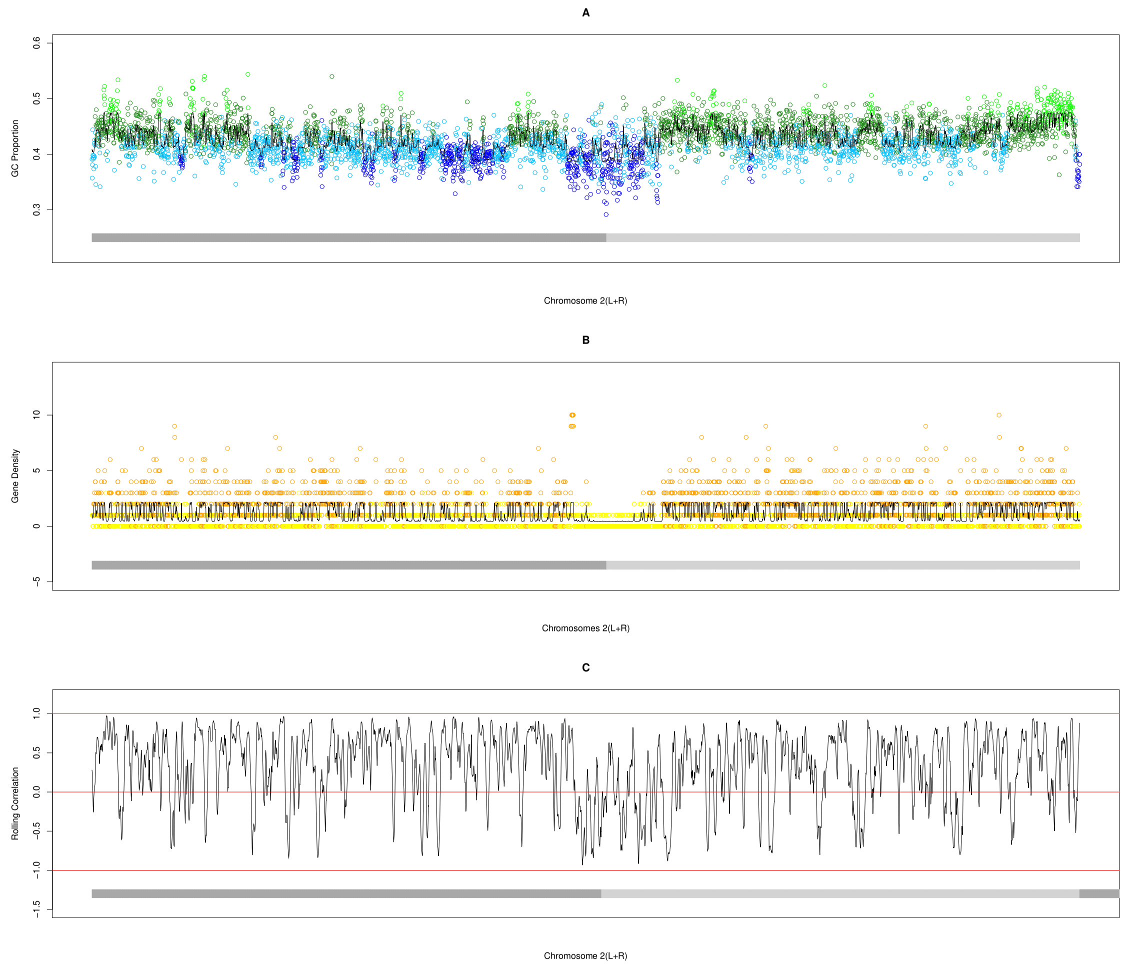


Figure 8: In each of the above panels A-C, the *Drosophila melanogaster* chromosome 2 (with the left and right demarked by dark and light grey horizontal bars respectively) are plotted for different oHMMed analyses: In A, the average GC proportion in every 10kb window is coloured by the oHMMed-inferred GC state. The two lower states are in blue and the two higher ones in green, with the shades lightening with higher GC proportion. In B, the number of protein coding genes for every 10kb window are shown in colours corresponding to the oHMMed-inferred gene density states: yellow and orange mark increasing gene density states. Note that in both A and B, the black lines trace the posterior (inferred) means returned by oHMMed with normal and gamma-poisson emissions respectively. These position-specific posterior means are the sum of estimated means times the respective probabilities of each state, thus combining both estimated mean values and the algorithm's certainty of the assigned state. In C, Spearman's correlation for the two posterior means in shown in rolling windows of 50 collated 20kb windows.
