## AdditionalFile3 for "Inference of Genomic Landscapes using Ordered Hidden Markov Models with Emission Densities (oHMMed)"

### 1. Comparing oHMMed to its Unordered Counterpart

The key feature of the oHMMed is its utilisation of **convex** emission densities, which then enables the ordering of hidden states. Recall that convexity is achieved by assuming a common standard deviation in the case of normal emissions, and a common shape parameter for the gamma-poisson emissions. Relaxing the convexity assumption yields a more general unordered HMM with emission densities which we will call unoHMMed. In this section, we first document the changes in the oHMMed algorithm required to implement unoHMMed, and then compare the performance of the methods. We do this on both simulated sequences that exhibit clear autocorrelation (*i.e.,* were generated by oHMMed), as well as the human data from the main text.

#### *1.1. Implementation of unoHMMed*

##### *1.1.1. Transition Matrix*

To update the transition matrix $\mathbf{T}$, the steps corresponding to the restriction of transitions and the maintenance of symmetry, *i.e.,* the sampling and the addition of auxiliary variables from the Dirichlet distribution from Additional File 1, Section A1 (precisely: Step 3 of the oHMMed iteration procedure), can simply be excluded for both normal and gamma-poisson emission densities.

##### *1.1.2. Normal Emissions*

In the case of normal emission densities, the main difference between the ordered and unordered algorithms concerns the way in which the means and variances are sampled. Now that each state has a different standard deviation, these must be individually updated (compare to Additional File 1, Section A1, Step 2 of the oHMMed iteration procedure). Define:

$$SS_i^{(t)} = \sum_l \mathbf{1}_{\theta_l^{(t)}=i} \big(y_l^{(t)} - \bar{y_i}^{(t)}\big)^2 . \tag{1}$$

The posterior standard deviations can then be sampled as follows, with posterior degrees of freedom $\nu_{pi} = 1 + L_i$ and posterior scale parameters using a vector of priors $\sigma_0$:

$$\sigma_{pi}^{2(t)} = \frac{SS_i^{(t)} + \sigma_{0i}^2 + (\mu_{oi} - \bar{y_i}^{(t)})^2 \frac{L_i^{(t)}}{L_i^{(t)}+1}}{L_i^{(t)} + 1} . \tag{2}$$

By contrast, the means of each state can sampled as described for the oHMMed algorithms, and simply are not sorted (compare to Additional File 1, Section A1 , Step 2 of the oHMMed iteration procedure).

##### *1.1.3. Gamma-Poisson Distribution*

In the case of the gamma-poisson emission densities, the sampling of both the $\alpha$ and $\beta$ parameters in the MCMC sampling scheme must be modified. Without assuming a common $\alpha$ but individual $\alpha_i$ per state instead, the $\beta_i$ must be sampled from a Gamma distribution with the following posterior shape parameters per state: $\delta_{pi} = \alpha_i L_i^{(t)} + \delta_0$ (compare Additional File 1, Section A2 under 'Updating $\beta_i$' ). Again, the resulting, newly sampled $\beta_i$ are then not sorted.

For the individual $\alpha_i$ themselves, the conditional posteriors utilized in the sampling procedure are proportional to (compare Additional File 1, Section A2 under 'Updating $\alpha$' ):

$$\Pr(\alpha_i \,|\, \boldsymbol{\theta}, \boldsymbol{y}, \beta_i^{(t+1)}) \propto \left( \frac{\upsilon_{pi}^{\alpha_i - 1}}{\Gamma(\alpha_i)} \right)^{\lambda_{pi}} , \tag{3}$$

with

$$\lambda_{pi} = \lambda_{0i} + L_i \, ,$$

where $\lambda_{0i}$ is a vector of priors and

$$\upsilon_{pi} = \left( \upsilon_0^{\lambda_{0i}} \prod_{l=1}^{L} \mathbf{1}_{y_l = i} \big( \beta_i^{(t+1)} y_l \big) \right)^{\frac{1}{\lambda_{0i} + L_i}} .$$

Given the new $\alpha_i^*$ sampled from the jump distribution as was described in the main text, the decision rule ratio is applied to each of these separately.

*1.1.4. Implementation Note*

Finally, note that unoHMMed as described here was implemented as an equivalent to oHMMed rather than a fully fleshed out method in its own right. Further modifications unique to unoHMMed (such as tuning of updates, or re-parameterising the models) may improve convergence behaviour. As such, the convergence traces generated by unoHMMed generally do not mix as well as those of oHMMed, particularly on long sequences and especially so for gamma-poisson emission densities. Situations in which mean and median log-likelihood of a fitted model differ drastically (for examples for oHMMed see e.g. $K = 6$ for the human GC proportion in the main text, or $K = 2$ for the human gene content in the main text, and the variance in convergence traces are also very large then become the norm rather than a rare occurrence indicative of poor model specification; this can be seen explicitly in the evaluation of the log-likelihood of unoHMMed with poisson-gamma emissions on simulated sequences Fig. 4B.

#### *1.2. Results*

##### *1.2.1. Summary and Interpretation*

The series of results presented in this subsection serves to emphasise that oHMMed was designed to infer a well-fitting model by assigning every one of a sequence of observations to a fixed number of hidden states with mean emission densities that can be discriminated with high statistical certainty. By contrast, unoHMMed appears to prioritise the model fit in terms of the overall emission density over clearly separated states. Recall, however, that the hidden states in this case are also not directly comparable using a single metric. For our simulations, we nonetheless sorted the states inferred by unoHMMed by mean emissions to facilitate the comparison of the methods. We chose to assess performance by collating the following results for each method across numerous inference runs on sequences that were simulated using the same underlying oHMMed model each time: the log-likelihood and the percentage of correctly assigned hidden states (which evaluate model fit), the deviation of the estimated values of the model parameters from the true values (which show general inference accuracy), and the number of inferred hidden states with significantly different mean emission densities. For the metrics of model fit and inference accuracy, the variances and means of the repetitions are compared between methods: F-tests are applied to check for difference in variances, and subsequently for the difference in mean either a t-test (when variances are equal) or Welch's test (when they are not) is employed. The numbers of distinctly different hidden states across repetitions are compared between methods via a $\chi^2$-test. We report the significant results (below Figs. 1, 2, 3, 4).—Overall, the main question here is: Given provable autocorrelation in an observed sequence, for *e.g.,* along a genome, which method is best suited for handling it?

Generally, the simulations show that both methods yield very similar and appreciably accurate results in the case of normal emission densities; this is even the case with very short sequences (Fig. 1A,C,D), although performance is naturally improved with sequence length (Fig. 2A,C,D). With short sequences, the effect of oHMMed preferentially maintaining distinguishable hidden states (Fig. 1C) at the cost of overestimating the transition rates between states as well as the means of the state-specific emission densities (Fig. 1D) is more pronounced. At the same time, unoHHMed indeed tends to infer hidden states that are not statistically distinguishable by their mean emissions (Fig. 1C). This is largely because the estimated standard deviations per state are less accurate (with some being over- and others underestimated; Fig. 1E).

In the case of gamma-poisson emission densities, performance of oHMMed and unoHMMed differs more markedly, with oHMMed clearly outperforming unoHMMed in terms of overall model fit (Figs. 3A,B and 4A,B), although this only becomes very significant for longer sequences. Similarly to before, oHMMed most often maintains the number of statistically distinct hidden states while unoHMMed does not, especially for short sequences (Figs. 3C, 4C). This difference is further reflected by slight underestimation of both $\alpha$ and the transition rates between hidden states by oHMMed, and overestimation of the $\beta_i$ by unoHMMed

(Figs. 3D, 4D).

The patterns concerning model fit and parameter estimation are noticeable in the comparison of oHMMed and unoHMMed on the hominid data (see Sec. 1.2.4). However, annotation of the genome according to both mean GC content and mean gene density yield very similar results between the methods since the full autosomal data (and therefore very long sequences) were used.

#### 1.2.2. *Simulations with Normal Emissions*

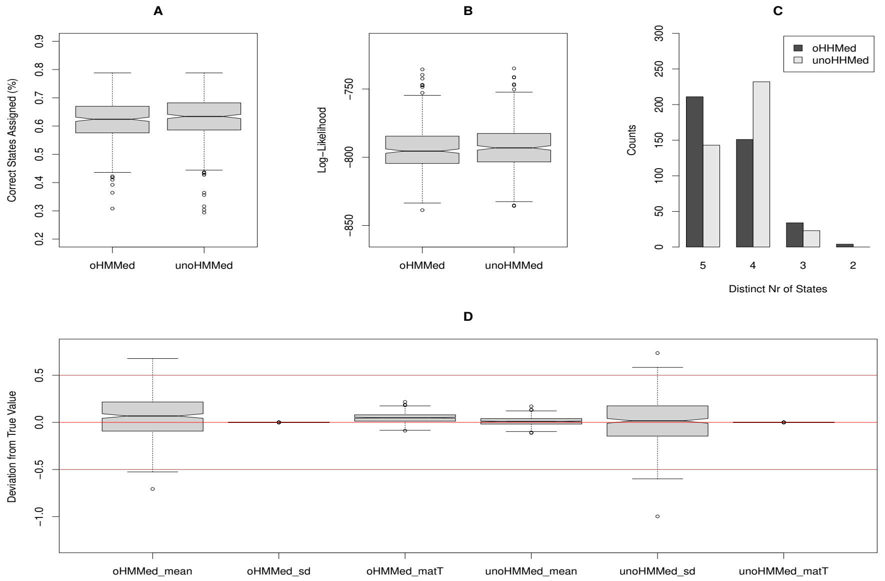


Figure 1: Performance of both methods was tested on sequences of length 500. These were simulated with oHMMed under the assumption of five hidden states emitting normal distributions with means of 1,2,3,4,5, a shared standard distribution of 1, and a probability of 0.1 of transitioning to any permitted neighbouring state. Then, both oHMMed and unoHMMed were used for inference; 2000 iterations (1000 of which were burn-in) were performed with prior and initial values as set per the usage recommendations for oHMMed on GitHub[1] . This sequence of simulation and inference was repeated 400 times, and the above figure evaluates the results across these repetitions. Note that although unoHMMed does not immediately allow ordering of hidden states, we sorted them by increasing mean emissions to facilitate comparison to oHMMed. In panel A and panel B, we show boxplots of the percentage of correctly assigned hidden states and the overall log-likelihood; means and variances of both measures are clearly non-significant between the two methods. Panel D shows barplots of how many of the mean emissions were deemed significantly different by one-sided t-tests (confidence level 0.95) after ordering the hidden states by increasing mean; the distribution of these counts differs significantly between methods ($\chi^2 = 36.311$, simulated p-val $< 0.05$). In Panel E, boxplots of the true values minus the inferred values for all the parameters are shown (averages in the cases of state-specific parameters). Between the two methods, the estimates of the mean differ significantly in both variance and means (F-val$= 21.363$ with df$= 1$, p-val$< 0.05$, and Welch t-val$= 4.329$ with df$= 436.27$, p-val$< 0.05$); the same is true for the estimated of the transition rates (F-val$= 2.173e^{31}$ with df$= 1$, p-val$< 0.05$, and Welch t-val$= 18.731$ with df$= 399$, p-val$< 0.05$). The estimates for the standard deviation differ only in variance (F-val$= 4.975e^{-33}$ with df$= 1$, p-val$< 0.05$).

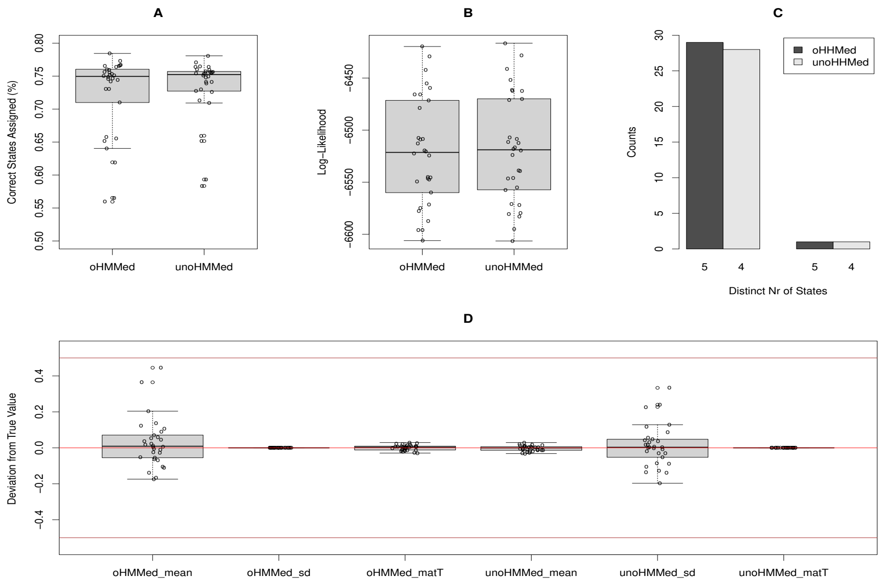


Figure 2: Keeping the same general set-up as in Fig. (1), we increased the sequence length to $2^{12}$ and performed 30 repetitions of the simulation and inference procedure. The above figure summarises the results across the repetitions. Since fewer of these were performed, the boxplots in panels A,B,E are superimposed on the individual values. The only significant difference in performance between the two methods lies in the variance of the estimators of the mean, standard deviation and transition rates (respectively: F-val$= 78.908$ with df$= 1$, p-val$< 0.05$; and F-val$= 1.443e^{-33}$ with df$= 1$, p-val$< 0.05$; and F-val$= 1.244e^{31}$ with df$= 1$, p-val$< 0.05$).

#### 1.2.3. *Simulations with Gamma-Poisson Emissions*

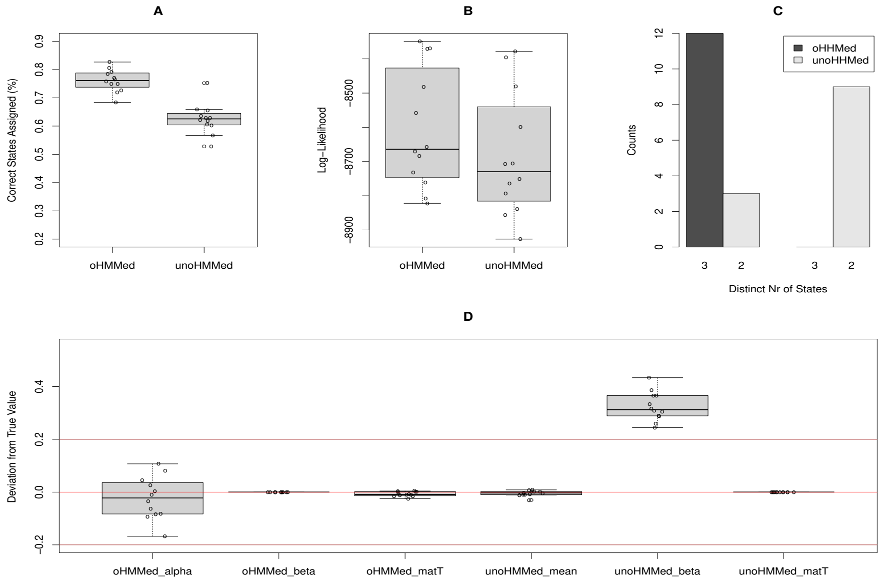


Figure 3: Performance of both methods was tested on sequences of length 2x$10^3$. These were simulated with oHMMed under the assumption of three hidden states emitting gamma-poisson distributions with shared $\alpha = 1.5$, respective $\beta_i$ of 0.029, 0.048, and a probability of 0.1 of transitioning to any permitted neighbouring state. Then, both oHMMed and unoHMMed were used for inference; 2000 iterations (1000 of which were burn-in) were performed with prior and initial values as set per the usage recommendations for oHMMed on GitHub[1]. This sequence of simulation and inference was repeated 12 times, and the above figure evaluates the results across these repetitions. Note that although unoHMMed does not immediately allow ordering of hidden states, we sorted them by increasing mean emissions to allow comparison to oHMMed. Note that since there were few repetitions of the testing procedure, the boxplots from panels A,B,E are superimposed on the individual values of the respective measure. In panel A and panel B, we show boxplots of the percentage of correctly assigned hidden states and the overall log-likelihood; the former measure differs significantly in mean between the two methods (Student's t-val=7.019 with df=22, p-val $< 0.05$). Panel C shows barplots of how many of the mean emissions of the ordered states were deemed significantly different by one-sided poisson rate tests with confidence level 0.95; the distribution of these counts differs significantly ($\chi^2 = 14.4$, simulated p-val $< 0.05$). In Panel D, boxplots of the true values minus the inferred values for all the parameters are shown (averages in the cases of state-specific parameters). Between the two methods, the estimates of $\beta$ differ significantly in both variance and mean (F-val$= 2.218e^{-32}$ with df$= 1$, p-val$< 0.05$, and Welch t-val$= -20.443$ with df$= 11$, p-val$< 0.05$); the same is true for the estimates of the transition rates (F-val$= 1.650e^{29}$ with df$= 1$, p-val$< 0.05$, and Welch t-val$= -2.639$ with df$= 11$, p-val$< 0.05$). The estimates of $\alpha$ differ only in variance (F-val$= 59.358$ with df$= 1$, p-val$< 0.05$).

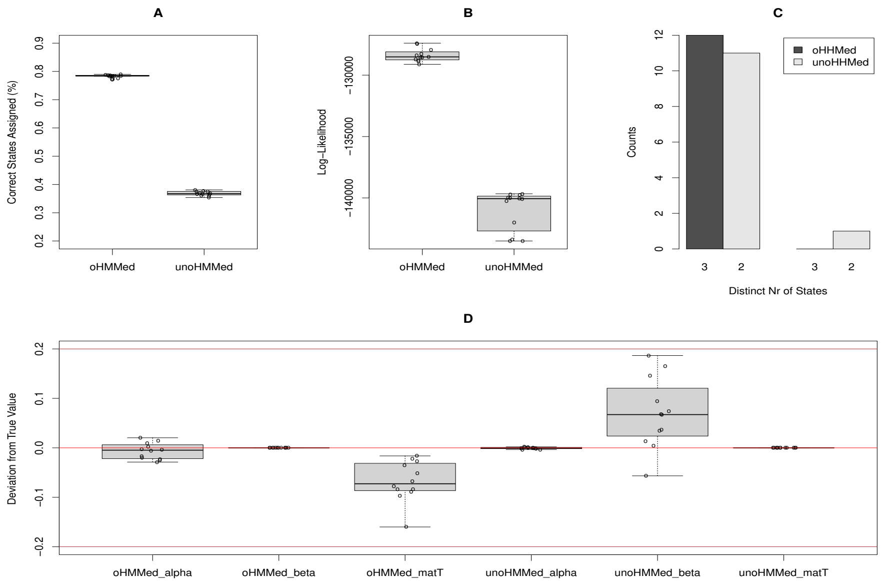


Figure 4: Here, we retained the set-up from Fig. (4) and increased the sequence length to $3\text{x}10^5$. We then performed 12 repetitions of the simulation and inference procedure. The above figure summarises the results across the repetitions. In panels A and B, one sees that the percentage of correctly assigned hidden states and the overall log-likelihood differ significantly in mean between the two methods, with oHMMed also being more precise in terms of the log-likelihood (respectively: Student's t-val= 149.9 with df= 22, p-val$<$ 0.05; and F-val= 0.112 with df= 1, p-val$<$ 0.05, and Welch t-val= 25.689 with df= 13.431, p-val$<$ 0.05 respectively). Further, panel D reveals that between the two methods, the estimates of $\beta$ differ significantly in both variance and mean (F-val= $7.138e^{-33}$ with df= 1, p-val$<$ 0.05, and Welch t-val= $-3.401$ with df= 11, p-val$<$ 0.05); the same is true for the estimated of the transition rates (F-val= $1.143e^{31}$ with df= 1, p-val$<$ 0.05, and Welch t-val= $-5776$ with df= 11, p-val$<$ 0.05). The estimates of $\alpha$ differ only in variance (F-val= 59.358 with df= 1, p-val$<$ 0.05).

#### *1.2.4. Hominid Data*

| GC Content (Normal Emissions) | oHMMed | unoHMMed |
|---|---|---|
| Sequence Length | 27067 | 27067 |
| Iterations (Burn-in) | 1500 (300) | 1500 (300) |
| Prior (=Initial) Means | 0.320 0.350 0.400 0.450 0.500 | 0.320 0.350 0.400 0.450 0.500 |
| Prior (=Initial) Standard Deviation | 1.1547 | 1.1547 1.1547 1.1547 1.1547 1.1547 |
| Prior (=Initial) Transition Rates | 0.990 0.010 0.000 0.000 0.000<br>0.010 0.980 0.010 0.000 0.000<br>0.000 0.010 0.980 0.010 0.000<br>0.000 0.000 0.010 0.980 0.010<br>0.000 0.000 0.000 0.010 0.990 | 0.990 0.010 0.000 0.000 0.000<br>0.010 0.980 0.010 0.000 0.000<br>0.00 0.010 0.980 0.010 0.000<br>0.000 0.000 0.010 0.980 0.010<br>0.000 0.000 0.000 0.010 0.990 |
| Log-Likelihood | 62315.69 | 64088.13 |
| Number of Distinct States | 5 | 5 |
| Estimated means | 0.3595 0.3912 0.4294 0.4832 0.5463 | 0.3573 0.3870 0.4226 0.4729 0.5356 |
| Estimated Standard Deviation | 0.0174 | 0.0120 0.0153 0.0199 0.0241 0.0285 |
| Estimated Transition Rates | 0.9612 0.0388 0.0000 0.0000 0.0000<br>0.0360 0.8828 0.0812 0.0000 0.0000<br>0.0000 0.1077 0.7774 0.1148 0.0000<br>0.0000 0.0000 0.2318 0.6806 0.0876<br>0.0000 0.0000 0.0000 0.2864 0.7136 | 0.9500 0.0478 0.0000 0.0000 0.0000<br>0.0360 0.8828 0.0812 0.0000 0.0000<br>0.0000 0.0908 0.7984 0.1108 0.0000<br>0.0000 0.0000 0.2099 0.6865 0.1036<br>0.0000 0.0000 0.0000 0.2826 0.7174 |

| Gene Density (Gamma-Poisson Emissions) | oHMMed | unoHMMed |
|---|---|---|
| Sequence Length | 28147 | 28147 |
| Iterations (Burn-in) | 40000 (5000) | 40000 (5000) |
| Prior (=Initial) Alphas | 0.9793 | 0.9793 0.9793 0.9793 |
| Prior (=Initial) Betas | 2.021 1.1355 0.2500 | 2.021 1.1355 0.2500 |
| Prior (=Initial) Means | 3.91711 0.8624 0.4846 | 3.91711 0.8624 0.4846 |
| Prior (=Initial) Transition Rates | 0.9238 0.0762 0.0000<br>0.2932 0.3529 0.3538<br>0.0000 0.8224 0.1776 | 0.9238 0.0762 0.0000<br>0.2932 0.3529 0.3538<br>0.0000 0.8224 0.1776 |
| Log-Likelihood | -26548.52 | -27201.28 |
| Number of Distinct States | 3 | 3 |
| Estimated Alphas | 5.9944 | 6.5815 7.3069 3.5681 |
| Estimated Betas | 44.5928 7.3796 2.0992 | 14.3396 10.6406 2.4022 |
| Estimated Means | 0.1346 0.8126 2.8565 | 0.2488 0.6867 2.7398 |
| Estimated Transition Rates | 0.9599 0.0.0401 0.000<br>0.0480 0.9381 0.0139<br>0.00 0.0622 0.9378 | 0.9076 0.0924 0.0000<br>0.1716 0.6011 0.2273<br>0.0000 0.0230 0.9770 |
