## AdditionalFile4 for "Inference of Genomic Landscapes using Ordered Hidden Markov Models with Emission Densities (oHMMed)"

### 1. Comparing oHMMed to its Unordered Counterpart (Continued)

In Supplementary Material, File 4 we extensively tested the performance of oHMMed vs its unordered counterpart, termed unoHMMed, on simulated sequences that conform to the modelling assumptions made by the oHMMed algorithms. This was intended to showcase the advantages of applying oHHMed when these assumptions hold on a given data set. To complement this, we here present the results of a smaller scale investigation of the performance of both oHHMed and unoHMMed on simulated sequences where the emission density is a mixture of normal distributions where each hidden state emits a single normal distribution component with a unique mean AND a unique standard deviation, and transitions between ALL hidden states are permitted. Obviously, the number of variable parameters is then greater than the number of parameters estimated by oHMMed. Therefore, it is unsurprising that inference with unoHMMed always yields a better fitting final model (see Fig. 1 for a comparison of the log-likelihoods, bearing in mind that a technically a more accurate comparison would be via maximum-likelihood and penalisation for differing number of parameters and that there is no consensus form for the penalisation in this case), and the individual estimates obtained by oHMMed will always be biased. Generally, the following pattern becomes clear: Since the oHMMed algorithm assumes that all hidden states share a standard deviation, it will infer a value near the average of all the true standard deviations. If this inferred value is high compared to the standard deviation of the components, oHMMed may effectively merge hidden states by inferring the same mean for them (see panels A in Figs. 2, 3). Otherwise, it will retain the correct number of hidden states and return slightly biased estimates of the emission densities. However, the sheer number of potentially confounded estimable parameter combinations leads to comparable behaviour in the unoHMMed algorithms: If the algorithm moves towards inferring an incorrectly high value for the standard deviation of a certain state, it may simultaneously push the mean for this state beyond the feasible parameter range and assign no or next to no observations to this state, rendering it redundant (see panels B in Figs. 2, 3). However, while inference of one fewer effective state in the case of oHMMed reduces the overall model fit (as evaluated by log-likelihood), a consequence of the increased flexibility of unoHMMed is that the same effect is not always observed (see Fig. 1). This reduction in the inferred number of hidden states actually occurs at a similar frequency for the oHMMed and unoHHMed algorithms respectively. Thus, while it is clear that unoHMMed has the potential to vastly outperform oHMMed on data sets that violate the model assumptions of the latter, inference may only be accurate with large data sets, a large number of iterations, and potentially a collation (aka averaging over the results) of multiple runs to take the failure of individual runs into account.

To illustrate these phenomena, we will show results pertaining to two separate scenarios: We chose to test the performance of both methods on sequences of length 12000, which we consider a moderate length. These were simulated with unoHMMed under the assumption of three hidden states emitting normal distributions with means of 1,2.5,5 respectively. In the first scenario, these states have very different standard deviations of 0.15 0.3 1.45 but a transition rate matrix that still only allows for transitions between neighbouring states (in order of increasing mean) specifically: $\begin{pmatrix} 0.9 & 0.1 & 0 \\ 0.1 & 0.8 & 0.1 \\ 0 & 0.1 & 0.9 \end{pmatrix}$.

In the second scenario, the true underlying standard deviations of the respective states are set to 0.15, 0.3, and 1.45 and the transition rate matrix is unconstrained, but still diagonally dominant: $\begin{pmatrix} 0.8 & 0.17 & 0.03 \\ 0.1 & 0.7 & 0.2 \\ 0.1 & 0.1 & 0.8 \end{pmatrix}$.

Then, oHMMed and unoHMMed were applied to the sequences from both scenarios 7000 iterations (5000 of which were burn-in) were performed with prior and initial values as set per the usage recommendations for oHMMed on GitHub (which means that in the case of unoHMMed, all prior and initial standard deviations are set to the same value). The process of simulation and inference was repeated 10 times for each value, and the figures on the next pages evaluate the results across these repetitions.

We note here that similar simulations with poisson-gamma emission densities were omitted, since the same general dynamics must hold but the results are less straightforward to interpret because of the less intuitive form of the emitted distribution.

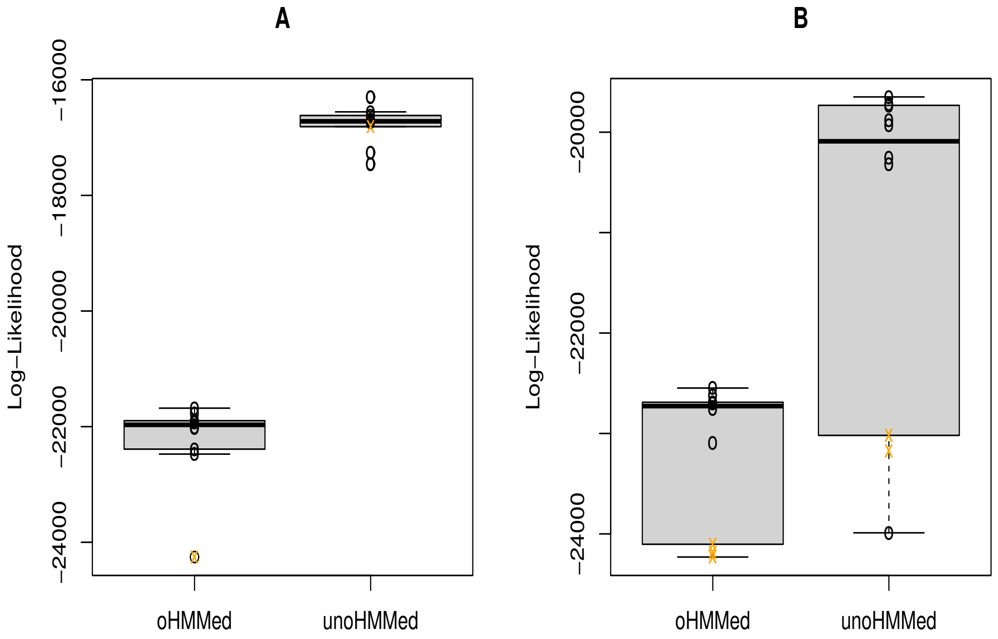


Figure 1: The overall log-likelihood of the final model inferred by the oHMMed and unoHMMed are shown in panel A for scenario 1 and panel B for scenario 2. The values of the individual runs that result in a reduction in the effective number of inferred hidden states, aka that lead to an incorrect inference of two hidden states rather than three, are marked with an orange 'x'. It is readily apparent that the estimates of the log-likelihood differ significantly in both means and variance between the two algorithms (F-val$= 5.0161$ with df$= 1$, p-value$< 0.05$, and Welch t-val$= -20.894$ with df$= 12.451$, p-value$< 0.05$ for scenario 1 and panel A, and F-val$= 0.17442$ with df$= 1$, p-value$< 0.05$, and Welch t-val$= -3.754$ with df$= 12.047$, p-value$< 0.05$ for scenario 2 and panel B).

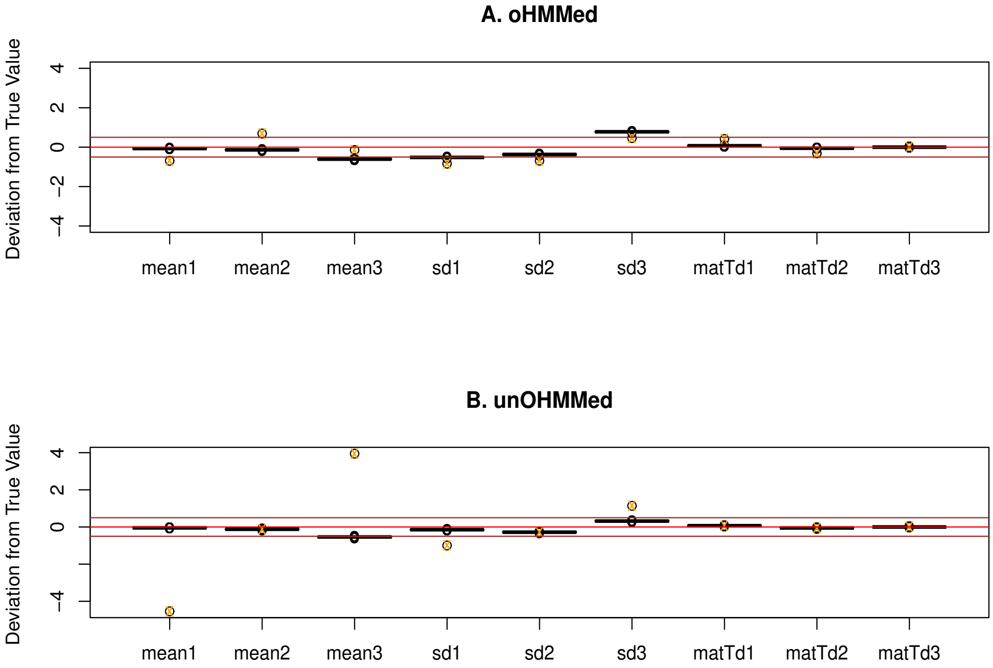


Figure 2: This plot shows the accuracy of the inferred parameters for the emission densities of the individual hidden states for scenario 1. The results obtained via the oHMMed algorithm are in panel A, and those obtained via the unoHMMed algorithm are in panel B. Both panels show, in order, the deviation of the inferred means from the respective true values, the deviation of the inferred standard deviation (for oHMMed) or standard deviations (for unoHMMed) from the respective true values, and the deviation of the average values of the main diagonal, first off- diagonal, and the corner entries in the inferred transition matrix from the respective true values. Once again, the values of the individual runs that result in a reduction in the effective number of inferred hidden states, aka that lead to an incorrect inference of two hidden states rather than three, are marked with an orange 'x'. Clearly, these runs lead to outliers in the above plots, and because of the different manner in which the algorithms achieve the effective reductions in states, the variance in estimation of values differs between methods. However, the only significant differences in mean estimation accuracy between oHMMed and unoHMMed occur for the standard deviation of the first and third hidden state (Welch t-val$= -3.5459$, with df$= 11.494$, p-value$< 0.05$ for the first, and Welch t-val$= 3.9487$ with df$= 11.653$, p-value$< 0.05$ for the third). This is expected since oHMMed infers only one standard deviation. Nonetheless, it is important to note the bias in estimating the means obtained from oHMMed and how it stems from the difference between the true standard deviations and the single estimate that oHHMed can infer.

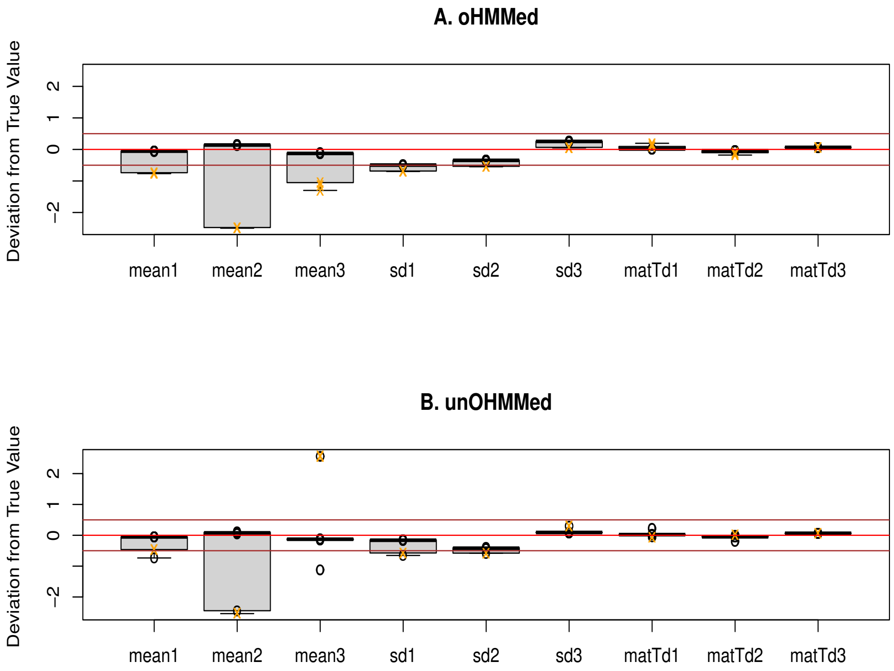


Figure 3: This plot shows the accuracy of the inferred parameters for the emission densities of the individual hidden states for scenario 2. The results obtained via the oHMMed algorithm are in panel A, and those obtained via the unoHMMed algorithm are in panel B. Both panels show, in order, the deviation of the inferred means from the respective true values, the deviation of the inferred standard deviation (for oHMMed) or standard deviations (for unoHMMed) from the respective true values, and the deviation of the average values of the main diagonal, first off- diagonal, and the corner entries in the inferred transition matrix from the respective true values. As always, the values of the individual runs that result in a reduction in the effective number of inferred hidden states, aka that lead to an incorrect inference of two hidden states rather than three, are marked with an orange 'x'. As before, these runs lead to outliers in the above plots; however, this time the resulting mis-estimation through these is comparable between methods in the sense that the variance in the inference accuracy is similar. Further, the only significant difference in mean estimation accuracy between oHMMed and unoHMMed here occurs for the standard deviation of the first hidden state (Welch t-val$= -3.5316$, with df$= 12.403$, p-value$< 0.05$)
