## AdditionalFile5 for "Inference of Genomic Landscapes using Ordered Hidden Markov Models with Emission Densities (oHMMed)"

### 1. Segmenting Human Chr1 By Epigenetic Markers

#### *1.1. Selecting the Number of Hidden States in oHMMed*

| EpiGeneticMark | Window Size (kb) | Number of Chosen States | Criteria |
|---|---|---|---|
| ATAC | 100 | n=4 | Log-Likelihood plateau roughly starts for n=4<br>the means of these states are distinct<br>and convergence behaviour is poor for n>4 |
| | 1 | n=5 | Log-Likelihood plateau clearly starts for n=5<br>the means of these states are distinct<br>and convergence behaviour gets worse for n>5 |
| H3K27ac | 100 | n=4 | Log-Likelihood plateau clearly starts for n=4<br>the means of these states are distinct<br>and convergence behaviour is poor for n>4 |
| | 1 | n=7 | Log-Likelihood plateau roughly starts for n=7<br>the means of these states are distinct<br>and the means for n=8 are no longer distinct |
| H3K27me3 | 100 | n=4 | Log-Likelihood plateau clearly starts for n=4<br>the means of these states are distinct<br>and convergence behaviour poor for n4 |
| | 1 | n=7 | Log-Likelihood plateau roughly starts for n=7<br>the means of these states are distinct<br>and increasing the number of states only further separates the 7th state |

Table 1: This table summarises the rationale behind how the most appropriate number of hidden states was chosen for each combination of epigenetic marker and window size; the selected number of hidden states is given in the third column and the reasoning in the fourth. For each data set consisting of counts of epigenetic marks per window, we ran oHHMed with between $n = 2$ and to up to $n = 9$ hidden states and considered the diagnostics for the output of each run according to our own usage recommendations on GitHub[1] in making the decisions.

### *1.2. Compilation of oHHMed Results for Each Epigenetic Marker*

Each table below summarises the most important results for each analysed epigenetic marker. For each of these, we provide the oHMMed-estimated parameters of the emitted poisson-gamma distributions in both 100kb as well as 1*kb* along chromosome 1 for the appropriately chosen number of hidden states. We further show what proportion of windows was assigned to each state, as well as the average number of consecutive windows per the same state that is expected in the data. Subsequent pages show plots of the data with overlayed oHMMed results (the inferred hidden states and the posterior inferred mean).

| **ATAC (Poisson-Gamma Emissions)** | **100kb** | **1kb** |
|---|---|---|
| Number of Distinct States | 4 | 5 |
| Estimated Alpha | 5.527 | 3.052 |
| Estimated Betas | 0.03148 0.02159 0.00386 0.00138 | 1.79132 1.75006 0.85715 0.09855 0.00655 |
| Estimated Means | 175.7 256.3 1434.4 4011.3 | 1.7040 1.7442 3.5612 30.980 465.91 |
| Estimated Variances | 5576.3 1186.0 371400.1 2897027.5 | 0.95126 0.99663 4.1547 31.436 7110.2 |
| Estimated Transition Rates | 0.863 0.137 0.000 0.000<br>0.140 0.441 0.418 0.000<br>0.000 0.411 0.460 0.129<br>0.000 0.000 0.530 0.470 | 0.008 0.992 0.000 0.000 0.000<br>0.864 0.069 0.067 0.000 0.000<br>0.000 0.076 0.852 0.072 0.000<br>0.000 0.000 0.474 0.332 0.194<br>0.000 0.000 0.000 0.619 0.381 |
| Prop. of Windows per State | 0.336 0.281 0.314 0.068 | 0.193 0.407 0.342 0.043 0.014 |
| Consecutive Windows per State | 9.737 1.738 1.911 1.883 | 1.000 1.960 8.933 1.490 1.625 |

Table 2: Results for ATAC

| **H3K27ac (Poisson-Gamma Emissions)** | **100kb** | **1kb** |
|---|---|---|
| Number of Distinct States | 4 | 7 |
| Estimated Alpha | 6.635 | 4.014 |
| Estimated Betas | 0.02373 0.01386 0.00448 0.00133 | 6.64091 1.17448 0.90545 0.47232 0.17884 0.04400 0.00977 |
| Estimated Means | 279.9 479.3 1484.3 4991.2 | 0.60622 3.4180 4.4335 8.4996 22.45 91.27 411.1 |
| Estimated Variances | 11794.5 34572.3 331475.4 3748289.1 | 0.09129 2.9102 4.8965 17.996 125.54 207.44 42091.7 |
| Estimated Transition Rates | 0.914 0.086 0.000 0.000<br>0.056 0.684 0.259 0.000<br>0.000 0.306 0.510 0.184<br>0.000 0.000 0.528 0.472 | 0.721 0.279 0.000 0.000 0.000 0.000 0.000<br>0.051 0.938 0.011 0.000 0.000 0.000 0.000<br>0.000 0.009 0.923 0.068 0.000 0.000 0.000<br>0.000 0.000 0.101 0.816 0.083 0.000 0.000<br>0.000 0.000 0.000 0.218 0.639 0.143 0.000<br>0.000 0.000 0.000 0.000 0.347 0.489 0.164<br>0.000 0.000 0.000 0.000 0.000 0.470 0.530 |
| Prop. of Windows per State | 0.228 0.372 0.299 0.101 | 0.039 0.288 0.345 0.206 0.078 0.033 0.012 |
| Consecutive Windows per State | 14.500 3.483 2.102 1.953 | 4.867 27.839 19.114 6.871 3.070 2.022 2.164 |

Table 3: Results for H3K27ac

| **H3K27me3 (Poisson-Gamma Emissions)** | **100kb** | **1kb** |
|---|---|---|
| Number of States | 4 | 7 |
| Estimated Alpha | 3.515 | 7.844 |
| Estimated Betas | 0.14202 0.01712 0.00306 0.00110 | 21.750 4.2349 1.3349 0.55438 0.27348 0.15496 0.08629 |
| Estimated Means | 33.838 236.62 1289.5 3194.1 | 0.36077 1.8528 5.8773 14.152 28.686 50.625 90.909 |
| Estimated Variances | 238.3 13821.2 421480.6 2896452.7 | 0.016587 0.43751 4.4027 25.527 104.89 326.70 1053.5 |
| Estimated Transition Rates | 0.425 0.575 0.000 0.000<br>0.053 0.552 0.395 0.000<br>0.000 0.166 0.622 0.212<br>0.000 0.000 0.143 0.857 | 0.947 0.053 0.000 0.000 0.000 0.000 0.000<br>0.069 0.822 0.108 0.000 0.000 0.000 0.000<br>0.000 0.098 0.772 0.131 0.000 0.000 0.000<br>0.000 0.000 0.113 0.740 0.146 0.000 0.000<br>0.000 0.000 0.000 0.102 0.768 0.130 0.000<br>0.000 0.000 0.000 0.000 0.134 0.812 0.054<br>0.000 0.000 0.000 0.000 0.000 0.241 0.759 |
| Prop. of Windows per State | 0.016 0.174 0.331 0.479 | 0.151 0.113 0.127 0.145 0.213 0.212 0.039 |
| Consecutive Windows per State | 1.929 2.474 2.772 7.900 | 23.701 6.826 5.284 4.645 5.427 7.022 4.811 |

Table 4: Results for H3K27me3

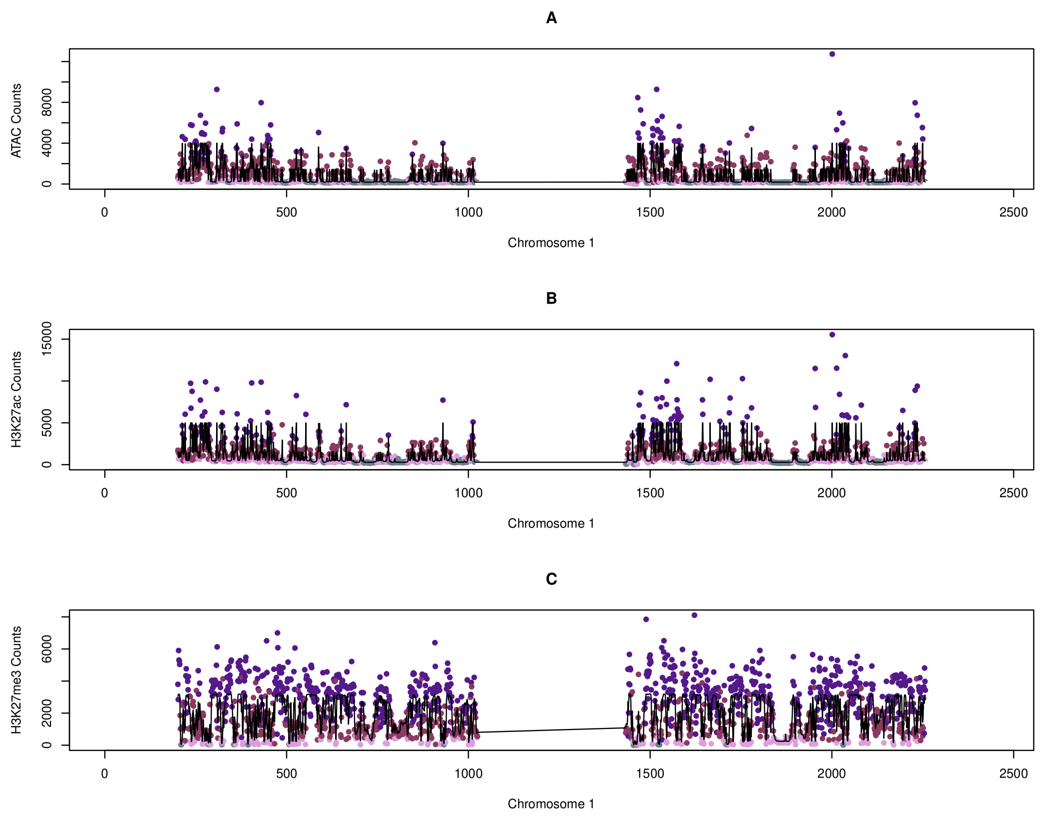


Figure 1: In each of the above panels A-C, the human chromosome 1 is plotted for different oHMMed analyses: In A, the number of ATAC counts per 100kb window are coloured by the oHMMed-inferred state the window has been assigned to. The state with the lowest average count is in grey, and then the states with increasing mean counts are in darkening shades of pink/purple. In panel B, the same plot is shown for H3K27ac, and in panel C, again the same plot for H3K27me3. The gaps on the sides and centre in all panels correspond to the telomere- and centromere-adjacent windows that were removed before the analyses. Note that in all panels, the black lines trace the posterior (inferred) means returned by oHMMed with gamma-poisson emissions. These position-specific posterior means are the sum of estimated means times the respective probabilities of each state, thus combining both estimated mean values and the algorithm's certainty of the assigned state.

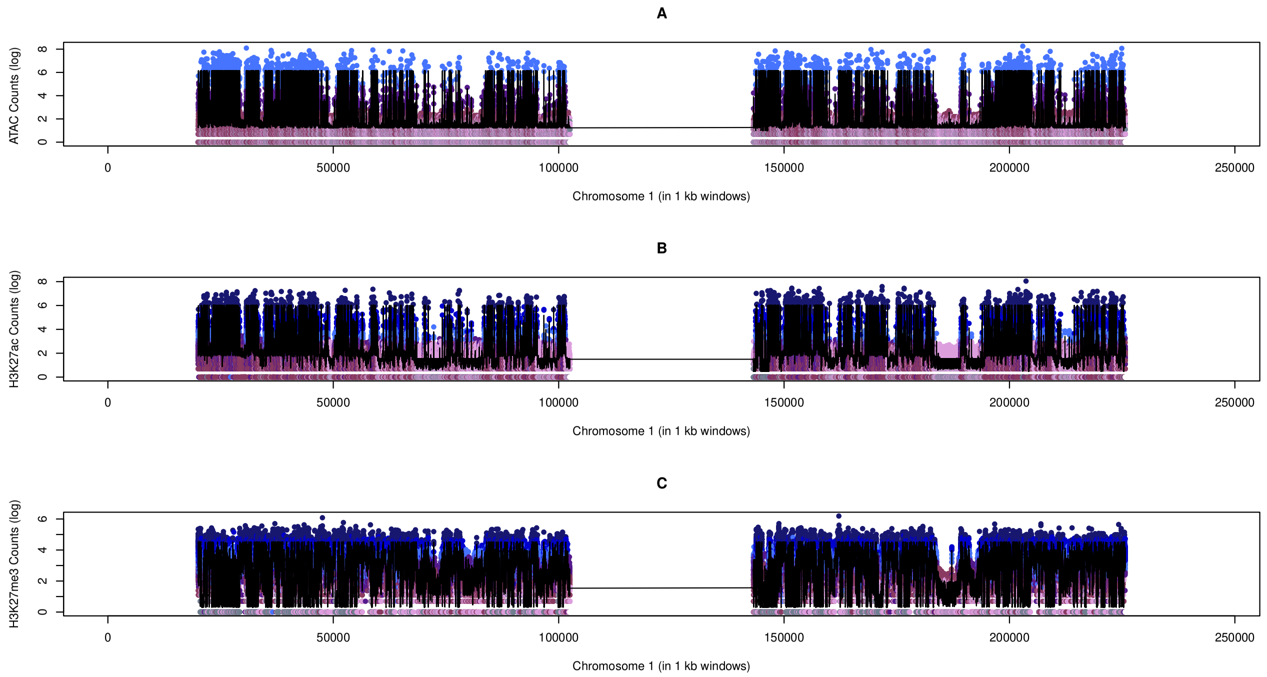


Figure 2: In each of the above panels A-C, the human chromosome 1 is plotted for different oHMMed analyses: In A, the number of ATAC counts per 1kb window are coloured by the oHMMed-inferred state the window has been assigned to. The state with the lowest average count is in grey, and then the states with increasing mean counts are first in darkening shades of pink/purple and then in darkening shades of blue. Importantly, note that the counts have been log-transformed in this case to increase visibility. In panel B, the same plot is shown for H3K27ac, and in panel C, again the same plot for H3K27me3. The gaps on the sides and centre in all panels correspond to the telomere- and centromere-adjacent windows that were removed before the analyses. Note that in all panels, the black lines trace the posterior (inferred) means returned by oHMMed with gamma-poisson emissions. These position-specific posterior means are the sum of estimated means times the respective probabilities of each state, thus combining both estimated mean values and the algorithm's certainty of the assigned state.
